# Autologous human iPSC-derived Alveolus-on-Chip reveals early pathological events of *M. tuberculosis* infection

**DOI:** 10.1101/2025.03.14.643230

**Authors:** Chak Hon Luk, Gabriel L. Conway, Kim Jee Goh, Antony Fearns, Irene Rodriguez-Hernandez, Nathan J. Day, Natalia Athanasiadi, Rocco D’Antuono, Enrica Pellegrino, Janick D. Stucki, Nina Hobi, Maximiliano G. Gutierrez

## Abstract

Immunocompetent and experimentally accessible alveolar systems to study human respiratory diseases are lacking. Here, we developed a single donor human induced pluripotent stem cell (iPSC)-derived Lung-on-Chip (iLoC) containing Type II and I alveolar epithelial cells, vascular endothelial cells, and macrophages in a microfluidic device that mimic lung 3D mechanical stretching and air-liquid interface. Imaging and scRNA-seq analysis revealed that the iLoC recapitulated cellular profiles present in the human distal lung. Infection of the iLoC with the human pathogen *Mycobacterium tuberculosis* (Mtb) showed that both macrophages and epithelial cells were infected and showed limited bacterial replication. Stochastically, large macrophage clusters containing necrotic core-like structure and Mtb replication were observed. A genetically engineered autophagy deficient iLoC revealed that after Mtb infection, macrophage necrosis was higher upon ATG14 deficiency without bacterial replication. Altogether, we report an autologous, genetically tractable human alveolar model to study lung diseases and therapies.

## Main

The human alveolus is an important tissue microenvironment for material exchange, while being safeguarded by tissue-resident immune cells against respiratory pathogens, including coronaviruses and *Mycobacterium tuberculosis* (Mtb)^1^. Given its significance in homeostasis^2^ and promise for drug delivery, many *in vitro* human models have been developed to circumvent the differences in anatomy, immune cell composition and disease pathogenesis between human and animals^3^. Motivated by the FDA Modernization Act 2.0^4^, organ-on-chip (OoC) technologies emerge as predictive tissue modeling tools and reliable alternative to animal testing backed by hypothesis-based and unbiased studies^5^. In the last decade, numerous efforts aimed to recreate pulmonary microenvironment on microfluidic devices (e.g., chips), where these systems can be categorized by their choices of organism (i.e. human or murine), cell source (i.e. cell line, primary cells or stem cell-derived) and target tissue (nasal, airway or alveolus). There are several reported multicellular airway-on-chips^6,7^ and alveolus-on-chips^8–17^ that are based on cell lines^14^ and primary cells^6–12,15–17^ or a combination of both^13^. A major challenge in building robust and scalable distal lung models is the access to a virtually unlimited source of reliable, standardized, phenotypically characterized, and immunocompetent alveolar cells from a tractable single donor. Despite the use of human induced pluripotent stem cell (iPSC) technologies satisfied most of these requirements and successful examples of developing alveolar epithelial cells^18–26^, work combining both OoC and iPSC to mimic the distal lung is limited.

Here, we report the engineering of a physiologically inspired human lung-on-chip that combines human iPSC with lung-on-chip technologies. We built an autologous, experimentally accessible, multicellular lung-on-chip from a single iPSC source that we named iLoC (for iPSC-derived <u>L</u>ung-<u>o</u>n-<u>C</u>hip) that enables incorporation of genetically engineered cells with an isogenic background. Using single cell transcriptomics, we demonstrate an important role of endothelial cells in regulating tissue innate and adaptive immune responses. Applying the iLoC as an early infection model for tuberculosis (TB), we found that both epithelial cells and macrophages are infected with Mtb, and cell death predominantly takes place in macrophages. Finally, by combining the iLoC with genetically engineered iPSC, we validated the role of the autophagy gene *ATG14* in regulating macrophage survival during Mtb infection.

## Results

### Lung progenitors differentiate into iAT2 and iAT1 cells in a lung-on-chip microfluidic device

To generate iPSC-derived type I and Type II alveolar epithelial cells (iAT1 and iAT2 respectively) and a biological barrier, we modified previously described protocols^20,22^ to differentiate iAT2 from human iPSC, passing through definitive endoderm (iDE), anterior foregut endoderm (iAFE), ventralized anterior foregut endoderm (ivAFE) stages to lung progenitors (iLungPro) that is enriched using a CPM^hi^ or CPM+ surrogate enrichment strategy (**Supplementary Fig. 1a**). This optimized protocol rendered the highest abundance of CPM+ iLungPro and required ventralization of anterior foregut endoderm compared to prior protocols^20,22^ (**Supplementary Fig. 1b-f**). We adopted the ^AX^Lung-on-Chip System, comprising of two electropneumatic devices, ^AX^Exchanger and ^AX^Breather that interfaces with the organ-on-chip device (AX12) to provide a physiological 3D stretching of the biocompatible silicon membrane as well as other maintenance function (Supplementary Fig. 1g)^8^. The enriched iLungPro were seeded directly on the Matrigel-coated AX12 to establish an epithelial layer both under static and breathing conditions (**Fig. 1a**), where mechanical stretch is introduced four days post-seeding as a strong trans-barrier electrical resistance (TER) developed (**Fig. 1b**). Recent works induced iAT2-iAT1 differentiation by pharmacological activation of YAP/TAZ pathway^18,26^. Herein, we initiated air-liquid interface (ALI) at seven days post-seeding to induce iAT2-iAT1 differentiation as it offers an accessible apical epithelium and minimizes medium composition change that is crucial to co-culture at later steps. Interestingly, the enrichment of CPM+ iLungPro was critical for barrier function (**Supplementary Fig. 1h**) and the TER was maintained in both static and breathing conditions, indicating the epithelial barrier remained functional up to 13 days (**Fig. 1b**).

**Fig. 1:**
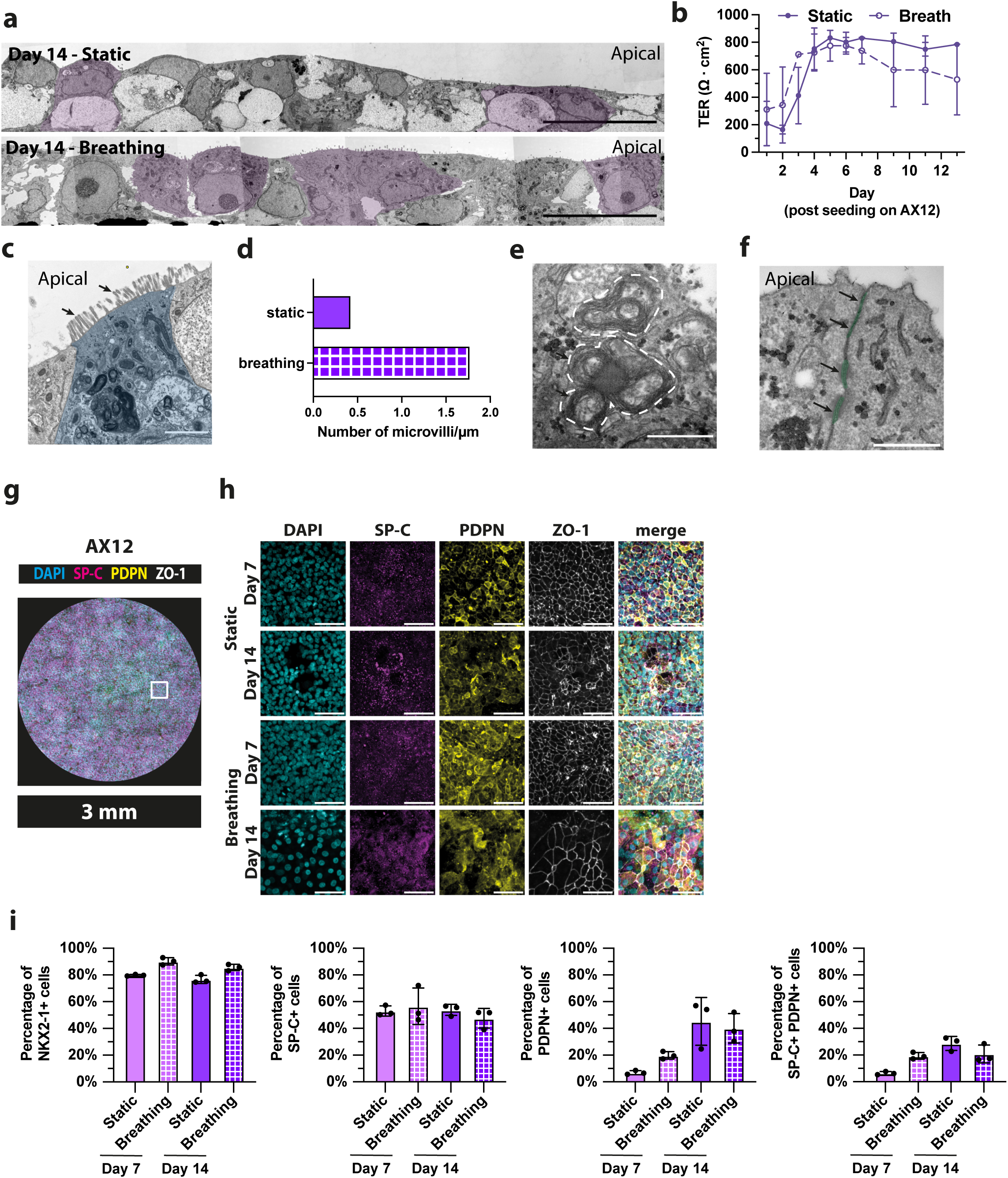
iPSC-derived iAT2 and iAT1s are differentiated in a lung-on-chip microfluidic device. **a**, Representative TEM images of iAT2 and iAT1 on AX12 at 14 days post-seeding, under static or breathing conditions. A fraction of the iAT2 with microvilli are highlighted in light purple. Scalebar = 20 µm. **b**, TER quantification of iAE on AX12 up to 14 days post-seeding, under static (solid line) or breathing (dotted line) conditions; mean ± s.d., n = 18 technical replicates from n = 3 independent experiments. **c**, TEM image showing a magnified view of microvilli on putative iAT2 cells (highlighted in blue) differentiated in AX12 at 21 days post-seeding, cilia are indicated with arrows. Scalebar, 2 µm. **d**, Quantification of microvilli ultrastructures in **c**, n = 30 quantified FOVs from 1 independent experiment. **e**, TEM image showing lamellar body-like structures in putative iAT2 cells differentiated in AX12 at 10 days post-seeding, lamellar bodies are highlighted by dotted lines. Scalebar, 500 nm. **f**, TEM image showing a magnified view of desmosomes in iAE differentiated in AX12 at 21 days post-seeding, desmosomes are highlighted in green and indicated with arrows. Scalebar, 500 nm. **g**, Representative image of the entire AX12 membrane with iAT2 and iAT1 cells showing nuclei (cyan), SP-C (magenta), PDPN (yellow) and ZO-1 tight junctions (white) at day 7 post-seeding, scalebar, 3 mm. **h**, Representative zoomed images of iAT2 and iAT1 differentiated on AX12 showing nuclei (cyan), SP-C (magenta), PDPN (yellow) and ZO-1 tight junctions (white) at day 7 and day 14 post-seeding, under static or breathing conditions. Equivalent area as depicted by square in **g**. Scalebar, 50 µm. **i**, Quantification of NKX2-1, SP-C and PDPN-expressing cells in iAE at day 7 and day 14 post-seeding, under static or breathing conditions; mean ± s.d., n = 3 independent experiments.

Using scanning electron microscopy (SEM), we observed the presence of microvilli structures covering the surface of the epithelium that prominently appeared at day 14 post-seeding (**Supplementary Fig. 2a**). Visualizing the microvilli structures using transmission electron microscopy (TEM) (**Fig. 1a and 1c**), we observed an enrichment of microvilli in the presence of mechanical stretch (**Fig. 1d**). At the ultrastructural level, TEM analysis showed that a substantial proportion of the cells exhibited typical microvilli features of AT2 cells and some of these cells displayed a cuboid morphology (**Fig. 1c**)^27^. Using fluorescence microscopy, we demonstrated the absence of tubulin-based ciliated structures at the apical side of iAT2, indicating the identity as microvilli (**Supplementary Fig. 2b**). Notably, we observed the presence of lamellar bodies (LB)-like ultrastructure with typically lamella structure and semi-dense core, organelles specific for storage and secretion of surfactants, in iAT2 cells^28^ (**Fig. 1e and Supplementary Fig. 2c)**. We also observed the presence of barrier function-implicated ultrastructure, tight junctions and desmosomes (**Fig. 1f**) as previously described in human lungs^29^. At day 14 post-seeding, we observed regions of different thicknesses and glycogen lakes^30^ formation contributed by the multilayers of cells, indicating the iPSC-derived alveolar epithelium (iAE) remains proliferative and harbor an intrinsic self-organization program (**Supplementary Fig. 2d**).

We phenotypically profiled the iAT2 and iAT1 differentiated on AX12 by single cell imaging analysis (**Fig. 1g**) and observed the expression of mature surfactant C (SP-C, iAT2 marker) and Podoplanin (PDPN, iAT1 marker) in differentiated iAT2 and iAT1 (**Fig. 1h**). At day 7 and 14, majority of the cells expressed the marker of distal lung epithelium NKX2-1 at high levels, where breathing motion achieved a higher number of NKX2-1+ cell (**Fig. 1i and Supplementary Fig. 3a**). Comparing the static and breathing conditions, we observed a comparable abundance of mature SP-C+ cells in both conditions at day 7 and 14, while a higher fraction of PDPN+ cells is observed under breathing condition at day 7 and both increased to a comparable fraction at day 14 (**Fig. 1h and i**). With dual labelling of mature SP-C and PDPN, we observed SP-C/PDPN double positive cells at day 7 and 14, indicating the presence of intermediate cell states. We also observed distinct morphologies of mature SP-C+ and PDPN+ cells (**Fig. 1h and Supplementary Fig. 3b**). We further validated the expression of iAT2 and iAT1 marker genes in the differentiated iAE and benchmarked to primary human adult lung samples. The levels of expression of *NKX2-1* (alveolar epithelial marker), *SFTPC*, *SLC34A2* (AT2 markers), *PDPN* and *CAV1* (AT1 markers) increased as iLungPro differentiated into iAT2 and iAT1, yet the overall levels were lower than that in human lung (**Supplementary Fig. 3c**). Altogether, we concluded that iAT2 and iAT1 differentiated on AX12 formed a functional epithelial barrier showing typical features of the alveolar epithelium observed in the adult human lung that includes the formation of lamellar bodies, cilia, tight junctions, and desmosomes.

### The microfluidic device enabled the formation of a functional epithelial-endothelial barrier

To differentiate vascular endothelial cells (iVEC) from human iPSC for the basal component of iLoC, we adopted a previously described protocol^31^. We validated the expression of endothelial marker, CD31 and vascular endothelial marker (**Supplementary Fig. 4a**), CD144 by fluorescence microscopy, cell surface expression of CD31, CD34, CD144 and KDR by flow cytometry (**Supplementary Fig. 4b**) as well as *CD31*, *CD34*, *CD144* and intracellular marker von Willebrand factor (*vWF*) by RT-qPCR against human umbilical vein endothelial cells (HUVEC) (**Supplementary Fig. 4c**). To confirm the cellular function of iVEC, we evaluated the capacity of very-low-density lipoprotein (vLDL) uptake and tubules formation in the angiogenesis assay (**Supplementary Fig. 4d** and e). The differentiated iVEC were seeded onto the basal side of the AX12 and iLungPro were subsequently seeded onto the apical side 1-2 days after the seeding of iVEC. The three cell types (iAT2, iAT1 and iVEC) together formed an epithelium-endothelium bilayer that maintained an ALI and TER up to 14 days under both static and breathing conditions initiated at the same time as iAE-only cultures (**Fig. 2a and b**). Importantly, the iAE and iVEC remained as intact bilayers and basal iVEC displayed a sustained vWF expression and a typical arrangement of actin cytoskeleton after 15 days of co-culture (**Fig 2c**). The patterned expression of mature SP-C and PDPN was maintained in the iAE despite different cell morphology to that in the absence of iVEC, and ZO-1 tight junctions were present in both iAE and iVEC endothelium (**Fig. 1h and 2d**). We concluded that the three cell types were maintained in the AX12 as an epithelial-endothelial functional barrier and preserving both epithelial and endothelial cellular phenotypes for at least 14 days.

**Fig. 2:**
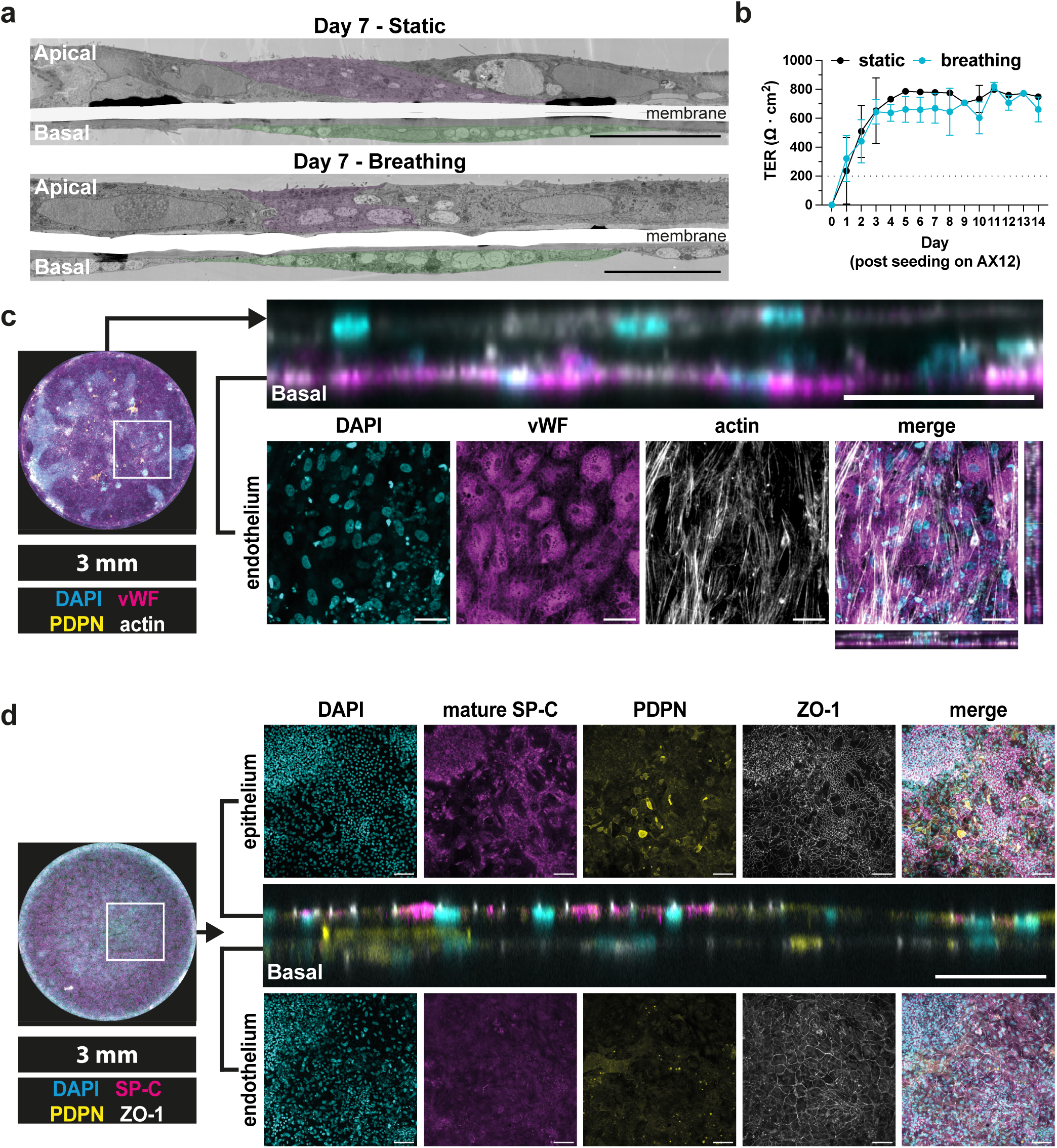
iPSC-derived vascular endothelial cells (iVEC) with iAE barrier build a dual epithelium/endothelium in a lung-on-chip microfluidic device. **a**, Representative TEM images of iAT2 and iVEC on AX12 at 7 days post-seeding, under static or breathing conditions, representative iAT2 and iVEC are highlighted in purple and green, respectively. Scalebar, 20 µm. **b**, TER measurement of iLoC up to 13 days post iLoC assembly under static (black) or breathing (cyan) conditions; mean ± s.d., n = 18 technical replicates from n = 3 independent experiments. (C) Representative images of orthogonal and planar views of iVEC cultured on basal side of AX12 together with apical iAT2 and iAT1 cells at day 15 post iLoC assembly, images showing nuclei (cyan), vWF (magenta) and actin (white). Scalebar, 50 µm. (D) Representative images and orthogonal and planar views of iVEC cultured on basal side of AX12 together with apical iAT2 and iAT1 cells at day 15 post iLoC assembly, images showing maximum projection of epithelium and endothelium, nuclei (cyan), mature SP-C (magenta), PDPN (yellow) and ZO-1 (white). Scalebar, 50 µm (orthogonal view) and 100 µm (planar view).

### Functional macrophages integrated into the epithelial barrier of the microfluidic device

Human alveoli contain alveolar macrophages essential for tissue homeostasis and immune surveillance, inspiring us to incorporate iPSC-derived macrophages (iPSDM) into the iAE-iVEC co-culture. We used a well characterized protocol for monocytes and macrophages^32^, where monocytes were subsequently differentiated with GM-CSF to yield macrophages of an alveolar-like phenotype^33,34^ that is also known as an M1/M2 hybrid phenotype in human macrophages^35^. iPSDM analyzed by flow cytometry and fluorescence microscopy were positive for CD11b, CD16, CD14 and negative for CD163 in contrary to iPSC-derived monocytes (iPSDMono) (**Supplementary Fig. 5a** and b). Differentiated iPSDM were seeded to achieve an optimal ratio of 1:10 to 1:20 (macrophage/epithelial cells) in the iLoC to represent that in human alveolar space^36^. At the ultrastructural level, iPSDM were loosely associated with the epithelium, where the iPSDM displayed intracellular vacuoles (**Fig. 3a**). With the presence of iPSDM, the barrier function of the iLoC remained intact (≥200 Ω·cm^2^), and the breathing of the iLoC achieved a more stabilized barrier function after 15 days of seeding. The TER in breathing condition was closer to physiological relevant ranges that is likely contributed by the breathing motions (initiated on day 4 of iLungPro seeding) in mimicking alveolar physiology (**Fig. 3b**). Notably, we found that groups of macrophages were associated with discrete regions on the iAE, and the association of macrophages induced an invagination of the epithelial but not endothelial barrier (**Fig. 3c-e**). We confirmed the ZO- 1 tight junctions were intact on both the epithelium and endothelium, iPSDM and iVEC were CD16 and vWF positive, respectively (**Fig. 3d-e**). Furthermore, iAT2 (SP-C, ABCA3) and iAT1 (AGER) markers are present among the iAE in iLoC, where SP-C is also detected in iPSDM, demonstrating the uptake of surfactant proteins, a key homeostatic function of alveolar macrophages^37^ (**Supplementary Fig. 6a-d**). The iPSDM were dwelling in different niches in the iLoC, either residing on the epithelium or integrated beneath the epithelium (**Fig. 3f**). When we measured the secretion of cytokines, we observed the secretion of many inflammatory and homeostatic cytokines reported to regulate local immune response and alveolar homeostasis, including IL-1β, TNF-α, IL-6, IL-8, IL-10 and IFN-γ^38^ (**Fig. 3g**). There was a polarized secretion of macrophage homing factors^39,40^ (M-CSF, GM-CSF, IL-9), antagonistic immunoregulators^41^ (IL-1α, IL-1Rα) and regeneration regulators^42^ (SDF-1α), further demonstrating the iLoC as a functionally organized epithelial-endothelial barrier (**Fig. 3g**). The lower level of M-CSF and GM-CSF detected in apical side can be accounted by the uptake by iPSDM for cellular homeostasis. Altogether, these data show that the iLoC is stable for at least 6 days with 4 cell types in the presence of both ALI and breathing.

**Fig. 3:**
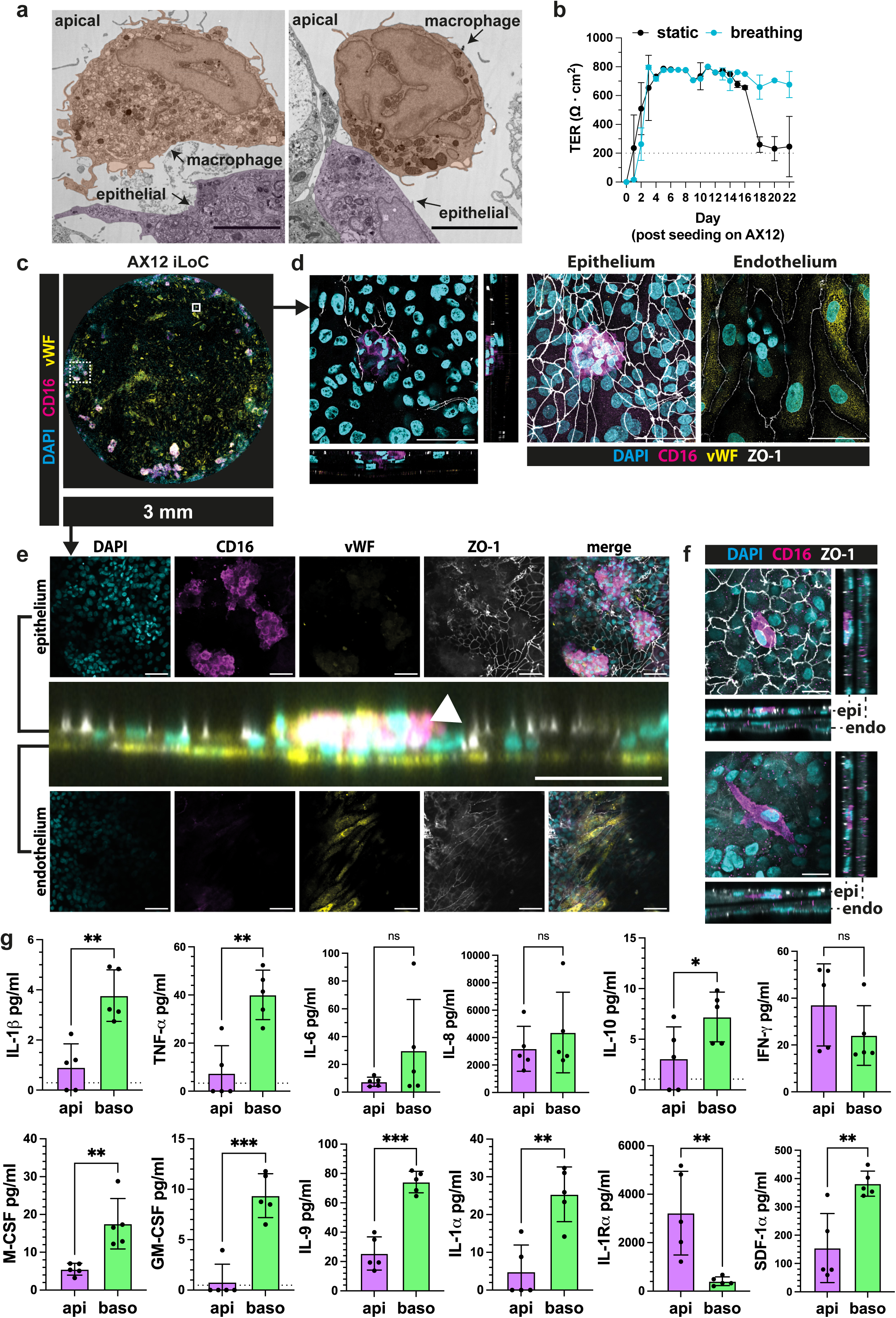
Incorporation of iPSC-derived macrophages (iPSDM) with iAE-iVEC barrier to form an immunocompetent alveolar microenvironment in a lung-on-chip microfluidic device. **a**, Representative TEM images of iLoC at day 15 post iLoC assembly without mechanical stretch, zoom in shows iPSDM resting on top of iAE. Representative iAE and iPSDM are highlighted in purple and orange, respectively. Scalebar, 5 µm. **b**, TER measurement of iLoC up to 22 days post iLoC assembly under static (black) or breathing (cyan) conditions; mean ± s.d., n = 18 technical replicates from n = 3 independent experiments. **C**, Representative images of an overview of AX12 iLoC images showing nuclei (cyan), CD16 (magenta), vWF (yellow) and ZO-1 (white). Scalebar, 3 mm. **d**, Representative confocal images depicted by solid line square in **c**, orthogonal and planar views of iPSDM residing on alveolar epithelium and maximum projection of epithelium and endothelium of AX12 iLoC, at day 21 post iLoC assembly, images showing nuclei (cyan), CD16 (magenta), vWF (yellow) and ZO-1 (white). Scalebar, 50 µm. **e**, Representative confocal images depicted by dotted line square in **c**, orthogonal and planar views of iLoC epithelium and endothelium with iPSDM (arrowhead) residing on alveolar epithelium, images showing nuclei (cyan), CD16 (magenta), vWF (yellow) and ZO-1 (white). Scalebar, 50 µm (orthogonal view) and 100 µm (planar view). **F**, Representative images of planar and orthogonal views of iPSDM residing on top of (top) and integrated beneath (bottom) alveolar epithelium at day 21 post iLoC assembly, images showing nuclei (cyan), CD16 (magenta) and ZO-1 (white). Scalebar, 20 µm. **g**, Quantification of IL-1β, TNF-α, IL-6, IL-8, IL-10, IFN-γ, M-CSF, GM-CSF, IL-9, IL-1α, IL-1Rα and SDF-1α secretion from the epithelial and endothelial side of iLoC; mean ± s.d., n = 5 independent experiments, Student’s t-test. p-value: ns P ≥ 0.05, * P < 0.05, ** P < 0.01, *** P < 0.001.

### The iLoC benchmarks human alveolar lung cells

The generation of the iLoC takes over 40 days from the iPSC stage, where macrophages are added and remain associated with the iLoC for at least 6 days (**Fig. 4a**). To define the cellular and phenotypic cell states on the iLoC, we profiled the iLoC using single cell RNA sequencing (scRNA-seq). We focused on the significance of endothelial cell and breathing motion to the iLoC, represented by co-culture of i<u>A</u>E-iPSD<u>M</u> (iAM) and iLoC under static or breathing conditions. Using a cellular marker classification approach, we identified the key cell types, iAT2, iAT1, proliferating iAT2 (pro. iAT2), iVEC and iPSDM alongside basal-like, ciliated-like and lung epithelial cells (iLE, **Fig. 4a and b, Supplementary Fig. 7a** and b). Using the defined cell type annotation, we detected an elevated expression of cell type-specific marker genes or a combination of the marker genes that contributed to the cell type annotations and tissue specificity^18,43^ (**Fig. 4c, Supplementary Fig. 7c**, 8a and b). It is notable that a couple of the key iAT2 and iAT1 markers are exhibiting a lower expression than recent literatures^18,25,26^ and primary alveolar cells, yet we are able to detect the other top ranked markers being reported^18^ (**Supplementary Fig. 7c**, 8a and c). Next, we mapped the sequenced cells onto a lung atlas with only datasets from healthy donors^43^ (**Supplementary Fig. 9a** and b). We observed the AX12-derived cells were mapped onto the clusters of epithelial cells, stromal cells, and macrophage/dendritic cells clusters (**Supplementary Fig. 9b**). Using clustifyr spearman correlation to examine gene expression similarity between annotated cell types in the lung atlas and our cluster cells, our cells were predicted to recapitulate a continuum of cellular states of alveolar epithelial cells as well as alveolar macrophages (**Supplementary Fig. 9c**). Combining the cellular marker annotation and cell type prediction, this analysis showed the successful recreation of an alveolar microenvironment that mimic endogenous cell-cell communication and intermediate cellular states.

**Fig. 4:**
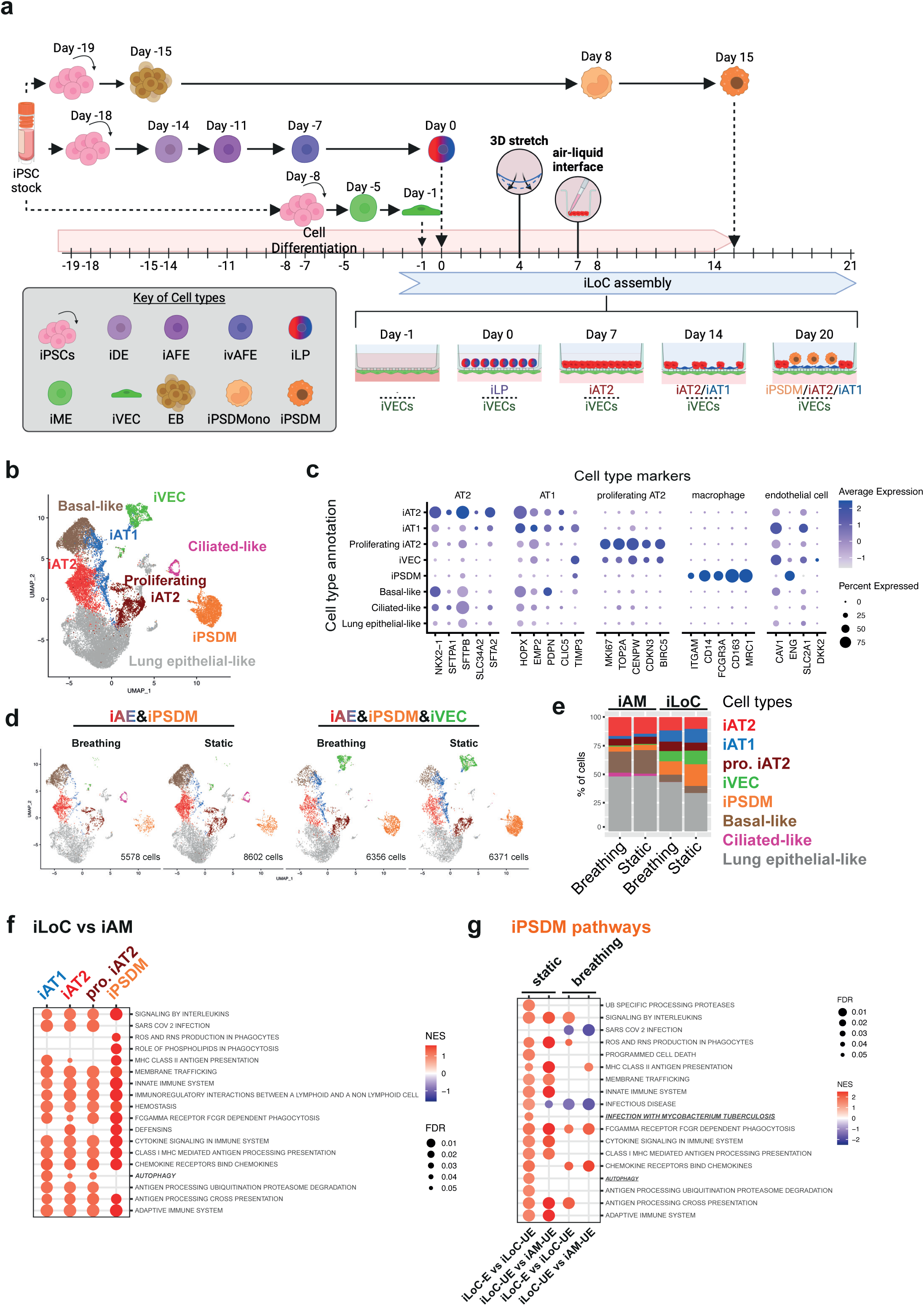
The four-cell type iLoC benchmarks human alveolar lung cells. **a**, Schematic outline of iAT2, iAT1, iVEC and iPSDM differentiation from iPSC and detailed steps of the iLoC assembly. **b**, Uniform Manifold Approximation and Projection (UMAP) with cell annotations of an integrated analysis of 26907 iAM or iLoC-derived cells in the presence or absence of mechanical stretch. (n = 1 per condition). **c**, Dot plot representation of the expression of selected AT1, AT2, proliferating AT2, macrophage and endothelial cell gene markers across annotated cell types. **d**, UMAP visualization of the integrated iAM or iLoC individual samples in the presence or absence of mechanical stretch from **b** split by experimental condition. **e**, Percentage of cells corresponding to the specific cell types (indicated with the color codes) in each individual sample from **b** and **d**. **f**, Dot plot showing selected Reactome enriched pathways from GSEA analysis for differential expression analysis in iAT2, iAT1, proliferating iAT2 and iPSDM cell types under static condition in the presence or absence of iVEC. (iLoC vs iAM) **g**, Dot plot showing selected Reactome enriched pathways from GSEA analysis for differential expression analysis between enriched and unenriched iPSDM subclusters in iLoC (iLoC-E vs iLoC-UE) and between unenriched iPSDM subclusters in iLoC and iAM (iLoC-UE vs iAM-UE) in presence or absence of iVEC and under breathing or static conditions.

The presence of an epithelial-endothelial barrier association and mechanical stretch by breathing are believed to be important components of the new generation lung-on-chip^5^. Thus, we experimentally analyzed the significance of breathing and presence of iVEC to iLoC using scRNA-seq. Comparing the four conditions, we observed a remarkable difference between the presence and absence of iVEC (iLoC vs iAM), where the presence of iVEC triggered a more stable iPSDM residence, iAT2-iAT1 trans-differentiation and distal lung cellular identity (**Fig. 4d and e**). Considering the annotated basal-like cells and iAT1 both expressed *PDPN*, this could account for the distinct morphology of PDPN+ cells in **Fig. 1h** and **2d**. On the contrary, breathing motion of the iLoC appears to have a less profound effect on the composition of cells (**Fig. 4d and 4e**). To better understand the impact of iVEC and breathing motion on iLoC cellular functions, we performed Reactome pathway analysis. We observed a general enrichment of immune-related pathways in iAT2, iAT1, pro. iAT2 and iPSDM in the presence of iVEC (iLoC condition). The enriched pathways in iAT2, iAT1 and pro. iAT2 included anti-viral and anti-bacterial pathways as well as oxidative phosphorylation; while iPSDM were poised towards lysosomal degradation, phagocytosis, cytokine responses and antigen presentation (**Fig. 4f**). Strikingly, the breathing motion induced a general down regulation of immune-related pathways in iAT2, iAT1, pro iAT2 and iPSDM while an up regulation in iVEC, indicating a cell type-specific impact of breathing motion and a potential role of mechanical forces in maintaining a naïve immune state (**Supplementary Fig. 10a and Supplementary Table 1**). Altogether, these data show that the iLoC benchmarked key cellular components of human lung.

### Basal endothelial cells impact apical macrophage and epithelial phenotypes in the iLoC

Owing to the significance of immune cells in alveolar homeostasis and combating foreign particles, we investigated the iPSDM phenotypes in the iLoC. For this, iPSDM cells were extracted, reanalyzed, and clustered *de novo*. We observed distinct subclusters uniquely enriched in the presence of iVEC (subclusters 2, 5, 6, 8, coined as iLoC-E) irrespective of the mechanical stretching, while the remaining are considered as unenriched clusters (subcluster 0, 1, 3, 4, 7, 9, coined iLoC-UE) (**Supplementary Fig. 11a-d**). The subclusters of iPSDM exhibited a high expression of *CD86* and *MRC1* as well as *CD68*, *TLR2*, *TLR4* (M1 markers) and *CD163*, *IL1R2*, *IL10RA* (M2 markers), aligning with previously reported M1/M2 phenotypes and cell type specific markers of alveolar macrophage and suggested the iLoC microenvironment represents that of alveolus for macrophage phenotype^35,44^ (**Supplementary Fig. 11e** and f). We observed one of the iPSDM subclusters was highly correlated to proliferating alveolar macrophages while some subclusters closer represented alveolar or interstitial macrophages (**Supplementary Fig. 11g**). Comparing the iLoC-enriched iPSDM subclusters to the iLoC-unenriched subclusters (iLoC-E vs iLoC-UE), the iLoC-enriched iPSDM displayed a higher expression in pathways of innate and adaptive immunity under both static and breathing conditions (**Fig. 4g**). At the same time, the unenriched iPSDM in iLoC (iLoC-UE) also upregulated these immune pathways as compared to the counterparts in iAM samples (iAM-UE), suggesting 3 populations of iPSDM of distinct expression of immune pathways (**Fig 4g**). Altogether, we demonstrated that presence of iVEC not only supported the homing of macrophages and global upregulation of immune pathways, but also increased the fraction of macrophages that upregulate key immune pathways. This indicates that co-culturing of macrophages with relevant cells in the iLoC environment imprints macrophage phenotype and function. Herein, we concluded that the iLoC mimics the cellular diversity and heterogeneity of human distal lung tissues.

### The iLoC mimics early stages of infection with *M. tuberculosis*

We next applied our iLoC system to model the alveolar tissue responses against the infection of *M. tuberculosis,* a human bacterial pathogen that targets the alveoli. We optimized the multiplicity of infection (MOI, number of bacteria per cell in the system) to 0.01 and aimed to use a very low bacterial number compared to conventional *in vitro* experimentation to achieve a closer mimicry to the physiological conditions of infection (e.g., as low as 2000 bacteria per iLoC). Upon Mtb infection from the apical side of iLoC, we observed infection of iPSDM and iAE by Mtb but not iVEC, where both iPSDM and iAE were infected (2.82±1.78% and 2.95±1.40%, respectively) (**Fig. 5a-d**). Considering the ratio of iPSDM to iAT2 and iAT1 in the iLoC, our findings agree with previous reports that infection predominantly takes place in macrophages in conjunction with other alveolar cell types^16,45^.

**Fig. 5:**
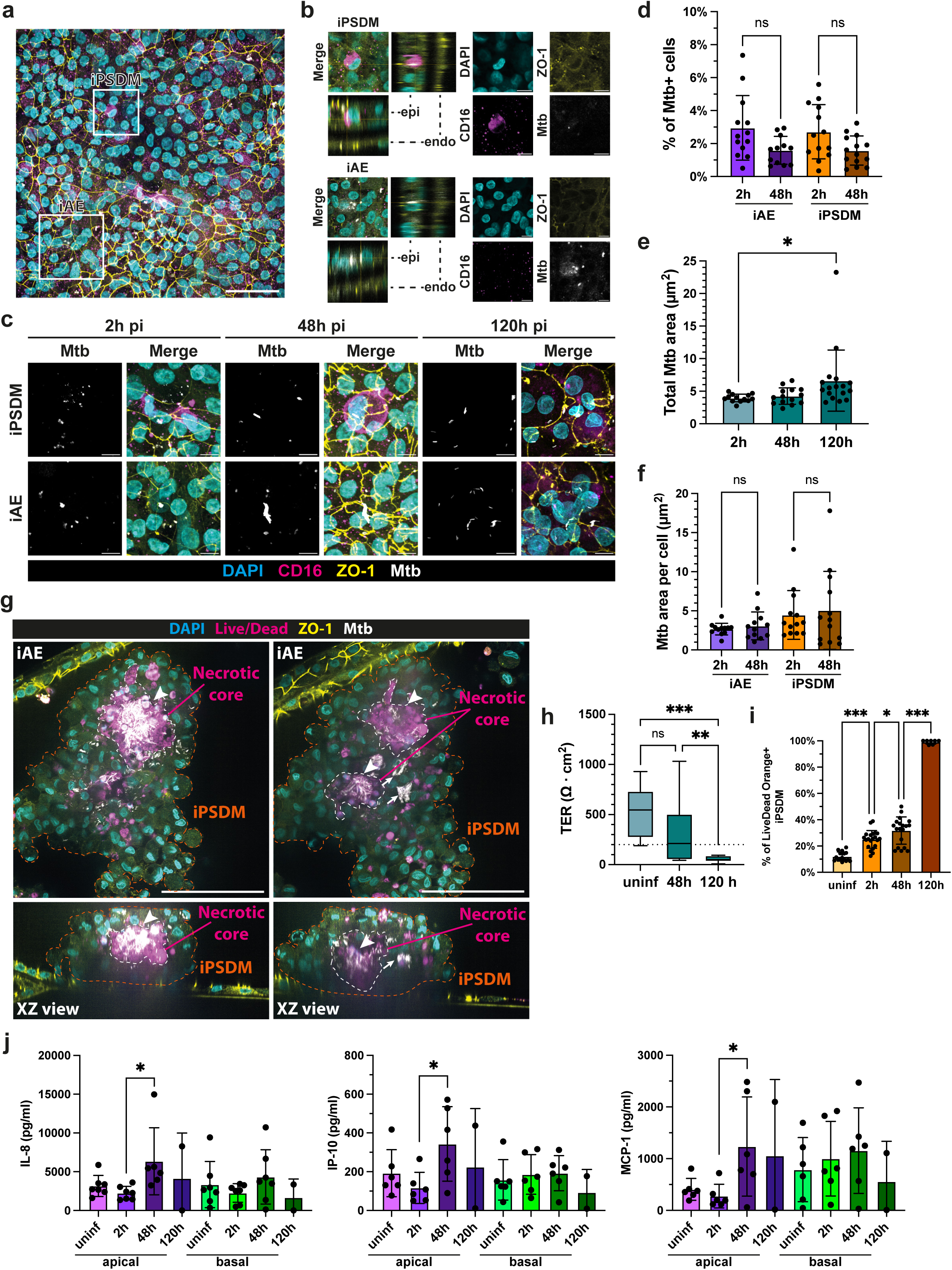
The iLoC mimic early stages of infection with *Mycobacterium tuberculosis*. **a**, Representative confocal image of Mtb-infected iLoC depicting Mtb dwelling in iAE, and iPSDM at 2 h pi. Scalebar, 50 µm. **b**, Zoomed images from **a** show nuclei (cyan), CD16 (magenta), ZO-1 (yellow) and Mtb (white). Scalebar, 10 µm. **c**, Representative images of Mtb replication in iAE and iPSDM at 2 h, 48 h and 120 h pi, images showing nuclei (cyan), CD16 (magenta), ZO-1 (yellow) and Mtb (white). Scalebar, 10 µm. **d**, Quantification of Mtb trophism in iAE and iPSDM at 2 h and 48 h; mean ± s.d., n = 12-13 analyzed FOVs from n = 3 independent experiments, one-way ANOVA. **e**, Quantification of Mtb area in iAE and iPSDM at 2 h and 48 h pi; mean ± s.d., n = 12-13 analyzed FOVs from n = 3 independent experiments, one-way ANOVA. **f**, Quantification of total Mtb area in iLoC at 2h, 48h and 120 h pi; mean ± s.d., n = 13-17 analyzed FOVs from n = 2-3 independent experiment, one-way ANOVA. **g**, Representative confocal and XZ orthogonal images from two separate z-planes showing the Mtb induced macrophage aggregation and formation of a necrotic core (positive for dead staining). **h**, TER measurement of iLoC uninfected vs infected with Mtb for 48 h; n = 6-14 technical replicates from n = 3 independent experiment, one-way ANOVA. **i**, Percentage of the iPSDM positive for Live/Dead Orange before and after Mtb infection; mean ± s.d., n = 7-21 analyzed FOVs from n = 3 independent experiment, one-way ANOVA. **j**, Quantification of IL-8, IP-10 and MCP-1 secretion from the epithelial and endothelial side of iLoC harboring iPSDM WT in uninfected, 2 h, 48 h and 120 h pi of Mtb infection; mean ± s.d., n = 2-6 independent experiment, one-way ANOVA. P-value: ns P ≥ 0.05, * P < 0.05, ** P < 0.01, *** P < 0.001.

In contrast to conventional 2D *in vitro* models, the majority of intracellular Mtb in both iPSDM and iAE did not significantly replicate within the first 48 h of infection (**Fig. 5f**). We then tested longer time points of infection. However, there was a disruption of the ZO-1 barrier in the iLoC at 120 h pi and the cell segmentation approach with ZO-1 used before (**Fig 5d, f**) for earlier infection timepoints was no longer suitable for reliable cell segmentation. To overcome this limitation, we applied another quantification approach that extracted the total Mtb area in iLoC, where we observed a significant increase in the Mtb area at 120 h pi (**Fig. 5f**).

We noticed the rare occurrence of large macrophage clusters containing necrotic core-like structure and Mtb replication (**Fig. 5g**). We detected a reduction but not a complete loss of barrier integrity as measured by TER, suggesting a tissue damage but not destruction of the iLoC at 48 h pi (**Fig. 5h**). As the infection progressed to 120 h pi, we observed sharp decline in TER accompanied with the destruction of both epithelial and endothelial barriers, indicating a collapse of the alveolar barrier function (**Fig 5h and Supplementary Fig. 12a**). This decrease in TER was associated with an increase in macrophage cell death as early as 2 h pi and subsequent epithelial cell death marked by a live/dead staining assay (**Fig. 5i, Supplementary Fig. 13a** and b). To evaluate the immune response elicited by the iLoC, we conducted a screen on the apical and basal cytokine expression and secretion of iLoC. With cytokine secretion profiling, we detected an elevation of IL-8 (48 h pi vs uninfected), IP-10 and MCP-1 (48 h pi vs 2 h pi) secretion on the apical but not the basal side of iLoC, indicating a polarized immune response against Mtb infection during the first 48 h of infection (**Fig. 5j**). We did not detect a further elevation of cytokine secretion level at 120 h pi, which could be attributed to the cell death and barrier disruption after 48 h of infection. Using RNAscope, we detected a rapid elevation of *IL-8* and *IL-6* expression in apical epithelium, which declined at later infection timepoints (**Supplementary Fig. 14a** and b). Altogether, we observed different kinetics of cytokine expression and secretion, highlighting the significance of capturing both readouts for spatiotemporal understanding of the underlying immune responses. Moreover, these data showed that the iLoC can recapitulate several aspects of Mtb infection and allowed us to define the early steps of tuberculosis and shed light on potential predictive cellular markers.

### A genetically engineered autophagy deficient iLoC reveals macrophage cell death without *M. tuberculosis* replication

We then took advantage of the autologous nature of our iLoC and incorporated genetically engineered cells to the iLoC. We have previously generated an *ATG14KO* iPSC line and derived iPSDM from it to demonstrate the role of ATG14 in Mtb infection in 2D *in vitro* models^46^. To define the role of ATG14 in macrophages in a physiologically relevant microenvironment, we generated a <u>G</u>enetically <u>E</u>ngineered iLoC (GE-iLoC) consisting of *ATG14KO* iPSDM and wild type iAT2, iAT1 and iVEC.

Upon Mtb challenge, we found both iPSDM and iAE were infected in both iLoC and GE-iLoC (**Fig. 6a**). Comparable levels of infection represented by the percentage of infected cells, total Mtb area and Mtb area per cell were detected in macrophages and epithelial cells of iLoC and GE-iLoC at 2 h pi (**Fig. 6b, 6c and d**). We detected a larger fraction of infected macrophages in GE-iLoC as compared to iLoC at 48 h pi, suggesting continuous bacterial uptake within the first 48 h of infection (**Fig. 6b**). Strikingly, total Mtb and Mtb loads in iAE and iPSDM in iLoC and GE-iLoC were similar at 2 h pi and 48 h pi, indicating macrophages were not permissive for Mtb replication within the first 48 h despite differences in bacterial uptake rate, as the infection progressed to 120 h pi, we observed similar levels of epithelial and endothelial barriers disruption and bacterial replication (**Fig 6c, d and Supplementary Fig. 10a**). Owing to the comparable pathological outcomes post-Mtb infection, there was no apparent paracrine effect on the antimicrobial capacity of epithelium when residing in an alveolar-like microenvironment. We further investigated the role of ATG14 in cell death in the iLoC^46^. Using a live/dead staining assay to evaluate the incidence of cell death in iLoC, another hallmark of early TB, we detected cell death prevalently in iPSDM within the first 48 h of infection, which was induced by Mtb infection as early as 2 h and *ATG14KO* iPSDM exhibited a higher basal as well as Mtb-induced cell death (**Fig. 6e and Supplementary Fig. 13a**). Both WT and *ATG14KO* iPSDM harbored comparable load of intracellular Mtb, while *ATG14KO* iPSDM exhibited an elevated cell death level, suggesting a bacterial load independent trigger of macrophage cell death (**Fig. 6d and e**). We observed a partial loss in TER at 48 h pi and further decline beyond 48 h pi in both iLoC and GE-iLoC, suggesting the loss of ATG14 in iPSDM does not affect barrier function decline (**Fig. 6f**). Despite the complete disruption of both epithelial and endothelial barriers at 120 h pi, we observed disruption of the endothelial barrier after 48 h of infection in the GE-iLoC, suggesting cell-cell communication between the infected apical cells with the basal endothelial barrier (**Supplementary Fig. 12a**). However, there was comparable cell division rate depicted by nuclear Ki67 labelling in the GE-iLoC as compared to iLoC at 48 and 120 h pi (**Supplementary Fig. 15a** and b), suggesting that ATG14 deficiency in the macrophages is associated with the GE-iLoC triggered disruption of the endothelial barrier without affecting proliferation. To further understand the influence of ATG14 deficient macrophages in the alveolar microenvironment, we profiled the iLoC apical and basal cytokines. We observed higher levels of apical G-CSF and M-CSF as well as a loss of polarized secretion of IL-1β in GE-iLoC, reflecting a remodeling of alveolar microenvironment by *ATG14KO* iPSDM (**Fig. 6g**). Upon Mtb infection, the GE-iLoC elicited similar IL-8 responses together with less prominent IP-10 and MCP-1 secretion, indicating a change in inflammatory response (**Fig. 6h**). Altogether, these data show that in an iLoC with autophagy deficient macrophages, infection with Mtb triggered higher macrophage cell death without prominent bacterial replication, where the macrophage-specific autophagy deficiency also led to the remodeling of alveolar microenvironment in both uninfected and Mtb-infected conditions.

**Fig. 6:**
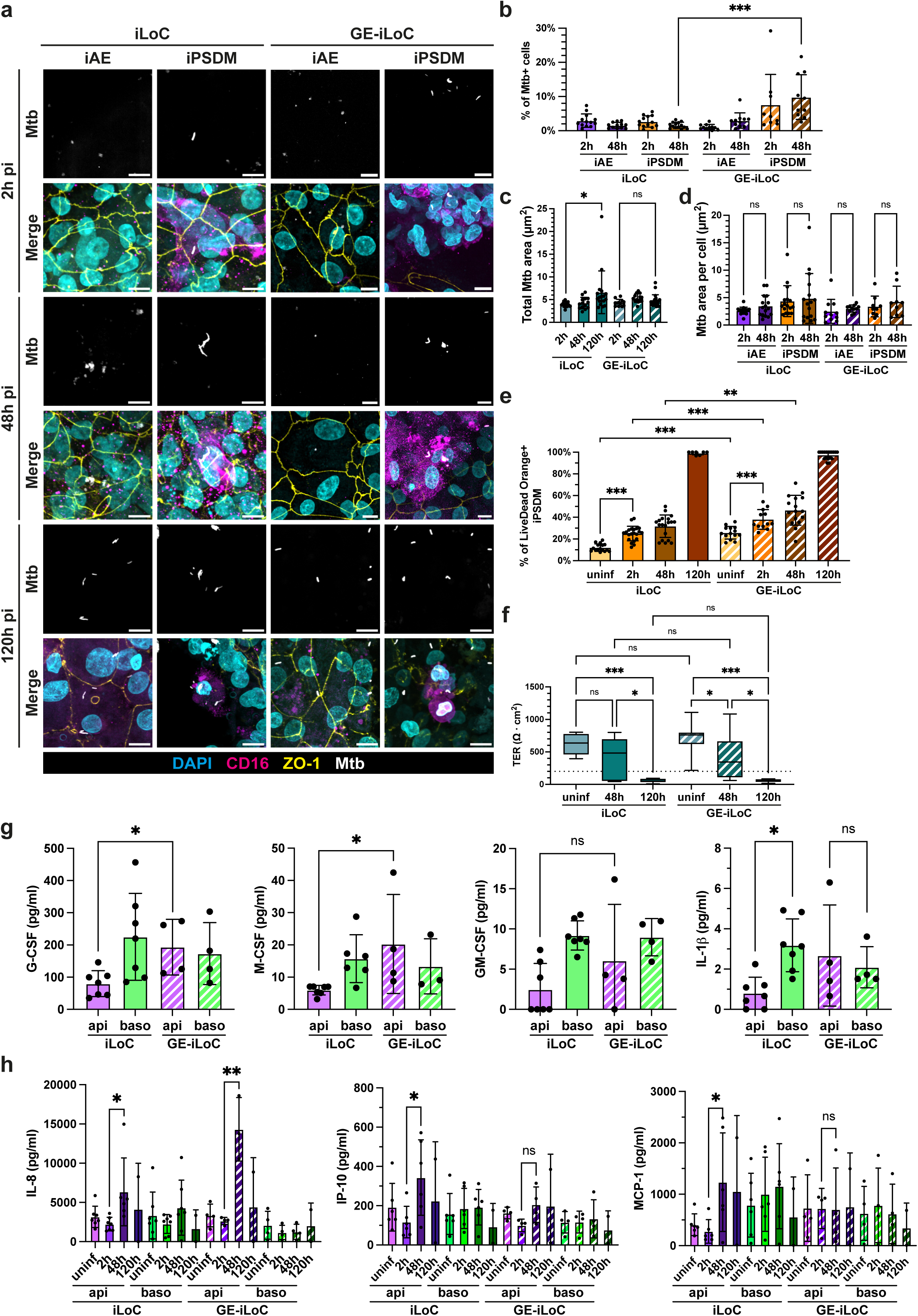
Mtb infection of cell type-specific genetic engineered iLoC shows induction of macrophage cell death without bacterial replication. **a,** Representative images of Mtb replication in iAE and iPSDM in iLoC or GE-iLoC at 2 h, 48 h and 120 h pi, images showing nuclei (cyan), CD16 (magenta), ZO-1 (yellow) and Mtb (white). Scalebar, 10 µm. **b**, Quantification of Mtb trophism in iAE and iPSDM in iLoC or GE-iLoC at 2 h and 48 h; mean ± s.d., n = 9-13 analyzed FOVs from n = 3 independent, one-way ANOVA. **c**, Quantification of total Mtb area in iLoC or GE-iLoC at 2h, 48h and 120 h pi in iLoC or GE-iLoC; mean ± s.d., n = 12-21 analyzed FOVs from n = 2-3 independent experiment, one-way ANOVA. **d**, Quantification of Mtb area in iAE and iPSDM at 2 h, 48 h pi; mean ± s.d., n = 9-17 analyzed FOVs from n = 3 independent experiment, one-way ANOVA. **e**, Quantification of Live/Dead Orange positive iPSDM WT and *ATG14KO* in uninfected, 2 h, 48 h and 120 h pi of Mtb infection; mean ± s.d., n = 7-21 analyzed FOVs from n = 2-3 independent experiment, one-way ANOVA. **f**, TER measurement of iLoC and GE-iLoC throughout the course of Mtb infection under static conditions; n = 6-14 technical replicates from n = 2-3 independent experiment, one-way ANOVA. **g**, Quantification of G-CSF, M-CSF, GM-CSF and IL-1β secretion from the epithelial and endothelial side of uninfected iLoC and GE-iLoC; mean ± s.d., n = 4-7 independent experiment, one-way ANOVA. **h**, Quantification of IL-8, IP-10 and MCP-1 secretion from the epithelial and endothelial side of iLoC and GE-iLoC in uninfected, 2 h, 48 h and 120h pi of Mtb infection; mean ± s.d., n = 2-7 independent experiment, one-way ANOVA. p-value: ns P ≥ 0.05, * P < 0.05, ** P < 0.01, *** P < 0.001.

## Discussion

To our best knowledge, this is the first report of a genetically engineered lung-on-chip that combines solely isogenic human iPSC-derived cells in an alveolar chip model^47^. We demonstrate the building of an autologous iLoC system comprising of Type I and II alveolar epithelial cells, vascular endothelial cells and alveolar macrophages with the capacity to incorporate CRISPR-Cas9 gene edited cellular components. Despite the successful recreation of an isogenic alveolar microenvironment, our system only represents a starting point for future improvements to the adoption of 3Rs principles (Replacement, Reduction and Refinement) and address the global demand to phase out animal models for drug efficacy and safety. Our chip system enabled mechanical stretching of the key alveolar cell types yet lacked fluidic flow, future efforts shall evaluate the impact of fluidic flow on our cellular components and incorporate additional cell types, e.g., fibroblasts and neutrophils to increase physiological relevance in tissue and immune modelling. Furthermore, the chip assembly is complex and time-sensitive due to the lack of long-term storage solution, cryopreservation approaches that offer good cell recovery and preservation of cellular phenotype will broaden the versatility of the system and scalability for applications in fundamental research as well as preclinical studies.

We harnessed the self-assembly capacity of iLungPro to generate different lung epithelial cell types without enrichment step subsequent to CPM+ iLungPro sorting. We generated a significant fraction of the intended alveolar epithelial cell types at an abundance comparable to the most recent reports in the field on cells before sorting^25^, while achieving a good specification to the lung lineage and the co-habitation of iAT2 and iAT1 including transitional and proliferating cell states. Such phenomenon suggests that our epithelial cells have retained their proliferative capacity and that the iLoC could represent a therapeutic assessment model of lung damage and regeneration. Considering the lower expression of key alveolar cell type markers as compared to the others^18,25,26^, future efforts are needed to address the long-term preservation of cell phenotypes in the iLoC. Using single cell transcriptomics, we captured the influences of mechanical stretching and endothelial signaling on the alveolar cells. Mechanical stretch induced a global pathway down regulation, which potentially implies a homeostatic state that offers a bigger dynamic range in tissue responses. Endothelial signaling appears to play diverse roles in the alveolar microenvironment, including promotion of iAT2 and iAT1 differentiation, macrophage home, global upregulation of inflammatory genes and antiviral pathways in epithelium and macrophages. Unexpectedly, the presence of endothelium leads to the enrichment of unique macrophage groups that exhibit stronger immune pathway up regulation, further highlighting the significance of endothelium in tissue immunity and the need of a multicellular system for physiological relevance.

Tuberculosis has a long asymptomatic phase that is challenging to be recapitulated by 2D cellular models and animal models due to the limited physiological resemblance^48^. Applying iLoC as early model of TB, we detected Mtb in macrophages and alveolar epithelial cells, aligning with previous reports of Mtb targeting not only macrophages^16,45^, where macrophages and epithelial cells are globally restrictive to Mtb replication as compared to 2D *in vitro* cultures, while stochastic replication is observed in conjunction with macrophage cell death up to two days post-infection. Such phenomenon can reflect the natural encounter of environmental Mtb, where humans are exposed to Mtb but only a small fraction of people developed TB with long asymptomatic phase and the underlying stochasticity remains largely unresolved. The iLoC displayed inflammatory cytokine secretion and barrier function deterioration in response to Mtb infection, reflecting the immuno-responsiveness of iLoC for early disease modeling. As the infection progressed to five days, we observed a rapid decline in barrier function, epithelium disruption and cell death. The observed phenomenon in iLoC is distinct from that in previous literature demonstrating alveolar macrophages being less restrictive than blood-derived or bone marrow-derived macrophages^44,49^. Such discrepancy could be attributed to differences in anatomical and genetic make-up between human and murine cells as well as the cell-cell interactions. Herein, the iLoC represents an alveolar system of strong inflammatory response that demonstrates tissue damage, further studies shall address the underlying molecular drivers and the impact on TB pathogenesis. Using a genetically engineered *ATG14KO* hybrid GE-iLoC, we found that ATG14 deficient macrophages are more susceptible to cell death under homeostatic and Mtb-infected conditions. We observed a higher bacterial uptake by macrophages from GE-iLoC, which has not been reported in macrophage 2D monoculture systems, and further investigation shall shed light on the pathophysiological implications of such phenomenon. Extracellular mycobacterial cording has been observed in other lung-on-chip models^15^ and we did not observe this predominately in our system, which could be attributed to multiple factors that differ between the two reported systems, including but not limited to cell sources, cellular physiology, microenvironmental cues and Mtb strains.

Respiratory infections and complications are among the top killers of human and major public health concerns. Fundamental understanding on the diseases will give us an upper hand against life-threatening diseases, where physiologically relevant model systems are vital. We demonstrated the building of iLoC and its capacity to study respiratory infection. This system can be applied to modelling and drug discovery of other pulmonary diseases as well as biosafety assessment. Nonetheless, the scalable and robust nature of our stem cell-based system enable the development of clinically relevant patient-derived systems to understand gene-drug interactions and develop personalized medicine.

## Material and Methods

### Human iPSC culture and maintenance

KOLF2HPSI0114i-kolf_2 (KOLF2) human iPSC was sourced from Public Health England Culture Collections (catalogue number 77650100). *ATG14* KO KOLF2 iPSC were generated as described before ^46^. To resume iPSC culture from cryopreserved stocks, 6 well tissue culture treated plates are coated with Vitronectin XF (STEMCELL Technologies), diluting 40 µl in 1 ml PBS (Thermo Fisher Scientific), incubated at room temperature for 1 hour), then the Vitronectin solution is replaced with 1 ml Essential 8 medium (hereafter E8, Thermo Fisher Scientific) + 10 µM Y-27623 (Tocris Bioscience) per well. Cryopreserved vials of iPSC were thawed in 37°C water bath and contents are transferred to 10 ml E8 in a 15 ml centrifuge tube. iPSC is collected by centrifugation at 300 x g for 3 minutes and plated in Vitronectin-coated plates in E8 + 10 µM Y-27632. From the following day onwards, the medium is changed to E8 without Y-27632 daily until confluent. To maintain iPSC culture, iPSC is washed with PBS and dissociated with Versene (Thermo Fisher Scientific, incubated at room temperature for 5 minutes). The Versene is then removed, and iPSC is resuspended to a suspension of small clumps in E8 or mTeSR Plus medium (STEMCELL Technologies). The iPSC is then split in a ratio of 1:6 into plates coated with Matrigel (Corning, according to manufacturer’s guide) in mTeSR Plus. iPSC cultures are passaged every 3-5 days using Gentle cell dissociation buffer (STEMCELL Technologies) or Versene.

#### Alveolar epithelial cells differentiation from human iPSC

iPSC were seeded at 2x10^6^ cells per well in Matrigel-coated 6-well plate in mTeSR Plus + 10 µM Y-27632. iPSC were cultured for 3 days as a monolayer in a chemically defined medium (STEMdiff definitive endoderm kit, STEMCELL Technologies) to induce the commitment of the cell population to the definitive endoderm state. Successful differentiation is indicated by a high incidence (>80%) of CXCR4+CKIT+ analyzed by flow cytometry. In the following differentiation steps, cells are cultured in base medium cSFDM^20^ (3:1 v/v mix of IMDM and DMEM/Ham’s medium F12 mix, 1% GlutaMAX, 0.5% N-2 Supplement, 1% B-27 Supplement, 0.05% BSA, 50 µg/ml ascorbic acid, 0.45 mM 1- Thioglycerol, 200 ng/ml primocin). Definitive endoderm cells are replated and cultured for 3 days in TGF-b/BMP4 inhibiting AFE medium (2 μM dorsomorphin, Selleck Chemicals, 10 μM SB431542, Selleck Chemicals) to commit to anterior foregut endoderm. Next, cells are cultured for 4 days in vAFE medium (20 ng/ml rhBMP4, Peprotech, 3.5 μM CHIR99021, Selleck Chemicals, 1 μM Retinoic acid, Merck) to achieve ventralization. Subsequently, the cells are cultured for 7 days in progenitor specifying LungPro medium (10 ng/ml recombinant human KGF, R&D Systems, Inc., 20 μM DAPT, Merck, 10 ng/ml FGF10, PeproTech, 3 μM CHIR99021) to derive NKX2-1+ lung progenitor cells.

Lung progenitors are dissociated using TrypLE (Thermo Fisher Scientific, 10-15 minutes at 37 °C), NKX2-1+ lung progenitor cells are enriched by fluorescence-activated cell sorting (FACS) using surrogate cell surface marker CPM. Enriched CPM+ lung progenitor cells are plated in Matrigel-coated AX12 plates at a density of 9.7x10^5^ cells/cm^2^ (a seeding density that enabled rapid coverage of AX12 membrane and iAE barrier formation) in Alveolar medium (10 ng/ml recombinant human KGF; 50 µM dexamethasone, Merck, 0.1 mM 8-bromoadenosine 3’:5’-cyclicmonophosphate, Merck, 0.1 mM 3- isobutyl-1-methylxanthine, Merck, 10 μM Y-27632) for 7+ days to yield alveolar epithelial cells. The apical medium is removed to form air-liquid interface to trigger iAT2 to iAT1 trans-differentiation ^24^.

#### Alveolar macrophage differentiation from human iPSC

For embryonic body (EB) formation using AggreWell 800 (STEMCELL Technologies), AggreWell is treated with Anti-Adherence rinsing solution (STEMCELL Technologies), centrifuged for 5 minutes at 300 x g and then washed with PBS and pre-incubated with E8 + 10 µM Y-27632. iPSC is harvested and plated onto AggreWell at 4x10^6^ cells per well with centrifugation at 100 x g for 6 minutes. For the following 3 days, the medium was changed daily to fresh EB medium (E8, 50 ng/ml BMP4; 50 ng/ml VEGF-A, PeproTech, #100-20; 20 ng/ml SCF, PeproTech, #300-07). To set up EB factories, EBs are harvested from AggreWell by pipette up and down and collected through a 40 µm cell strainer and seeded into tissue culture-treated flask in factory medium (X-VIVO 15, Lonza, 2 mM GlutaMAX; 50 µM β-Mercaptoethanol, Gibco, 100 ng/ml M-CSF, PeproTech, 25 ng/ml IL-3, PeproTech). For factory maintenance, the factory is fed weekly with factory medium. The factories begin to produce monocytes 4-5 weeks post-establishment. For macrophage differentiation, monocytes are collected from the factory supernatant by centrifugation at 300 x g for 5 minutes and plated into 10 cm dishes at 5x10^6^ cells per dish in differentiation media (X-VIVO 15, 2 mM GlutaMAX, 50 ng/ml GM-CSF, PeproTech). The seeded cells are boosted with fresh differentiation medium 3- or 4- days post seeding. Depending on the demand of macrophages, this process can be scaled up or down ^32^.

#### Endothelial cell differentiation from human iPSC

iPSCs were seeded at 700,000 cells per plate in Matrigel-coated 10 cm dish in mTeSR Plus + 10 µM Y-27632 and the medium is replaced with mTeSR Plus on the following day. iPSC were cultured for 3 days as a monolayer in a chemically defined medium N2B27 (50% DMEM/F12 medium, Gibco, 50% Neurobasal medium, Life Technologies, 2X B-27 Supplement, 1X N-2 Supplement, 0.1% β-Mercaptoethanol, 8 µM CHIR99021, 25 ng/ml rhBMP4) to induce the commitment towards lateral mesodermal state. The cells are then cultured for 2 days in VEC medium (StemPro-34 SFM, Life Technologies, 1X GlutaMAX, 200 ng/ml VEGF-A, 2 µm forskolin, Abcam) to achieve differentiation into endothelial cells. The endothelial cells are harvested and CD144+ endothelial cells are enriched using magnetic cell sorting.

#### Multicellular iLoC assembly

A complete diagram of the assembly of the iLoC is shown in Fig. 4a. Step 1: iVEC sorting and seeding. Differentiated cells were detached, magnetically enriched for CD144+ iVECs and counted according to “Endothelial cell differentiation from iPSC” and “iVEC enrichment by magnetic cell sorting” protocols. 8x10^4^ iVEC are seeded in 15 μl iVEC medium per membrane on basolateral side of AX12. iVECs are allowed to attach overnight (under unclosed condition) at 37 °C, 5% CO_2_. On the next day, the AX12 plate is initialized with iVEC medium (StemPro-34 SFM; 50 ng/ml VEGF-A; 10 µM Y- 27632).

Step 2: iLungPro sorting and seeding. Differentiated cells are detached, enriched for CPM+ iLungPro according to “Alveolar epithelial cell differentiation from iPSC” and “iLungPro enrichment by fluorescence-activated cell sorting” protocol. 8x10^4^ iLungPro are seeded in 70 μl Alveolar medium per membrane on the apical side of AX12. iLungPro are allowed to attach overnight at 37 °C, 5% CO2.

Step 3: Seeding of iPSDM. To detach differentiated iPSDM, cells are incubated with Versene, lifted using a lifter, collected by centrifugation and counted. To achieve a concentrated cell suspension for iPSDM seeding (2x10^6^ iPSDM/ml), desired amount of iPSDM is collected by centrifugation and resuspended in small volume of iLoC medium to 2x10^6^ iPSDM/ml. 10 µl of iPSDM cell suspension is added to each AX12 membrane to achieve an iPSDM:iAT2 & iAT1 ratio of 1:5-1:10.

We have generated iLoC using this protocol using three iPSC lines to date, including WTSIi018-B (RRID:CVCL_AE29), CRICKi018-A (RRID:CVCL_D0DD) and CRICKi019-A (RRID:CVCL_D0DE). In generating the figures of this work, we used the iPSC line WTSIi018-B, derived from a healthy donor (data not shown).

#### Maintenance of the iLoC

From day 2 onwards, apical and basal media are changed to fresh Alveolar medium and iVEC medium, respectively, every two days until the formation of air-liquid interface. Typically, a TER reading of ≥ 200 Ω·cm^2^ is detectable on Day 4 post seeding of iLungPro. To initiate 3D breathing after reaching a TER reading ≥ 200 Ω·cm^2^, the AX12 plate is placed into the ^AX^Dock connected to the ^AX^Breather (all components provided by AlveoliX), closed tight and 3D breathing of the desired chip is initiated. To form air-liquid interface, the apical medium is removed, and the basal medium is switched to iLoC medium (50% Alveolar medium; 50% iVEC medium) in a reduced volume to balance the hydrostatic pressure between the wells according to manufacturer’s instructions. After air-liquid interface formation, the basal medium is changed to fresh iLoC medium every two days. A detailed description on the design and handling of ^AX^Lung-on-Chip System has been reported recently ^14^.

#### Immunostaining and microscopy

For cells seeded on glass coverslips, cells were washed with PBS and fixed with 4% PFA (diluted from 16% stock, Electron Microscopy Sciences) in PBS for 15 minutes at room temperature. For cells seeded on AX12, half of the apical medium was removed and equal volume of 8% PFA in PBS is added; 4% PFA in PBS is flowed into the basolateral chamber to fix the cells for 15 minutes at room temperature.

For all samples infected with Mtb, samples were fixed in 4% PFA in PBS (final concentration) at 4 °C overnight to ensure complete killing of Mtb. The fixed cells were then washed 3 times with PBS and stored in 1% BSA PBS at 4 °C.

Cells were permeabilized with 0.01% saponin (Merck) in 1%BSA (Cell Signaling Technology) PBS for 1 hour at room temperature, then blocked in 3% BSA PBS for 1 hour. Cells were washed 3 times with PBS, then stained with primary antibodies in 1% BSA PBS for 1 hour at room temperature. Cells were washed 3 times with PBS, then stained with 4’,6-diamidino-2-phenylindole (DAPI, Invitrogen) and secondary antibodies in 1% BSA PBS for 1 hour at room temperature. Cells are then washed 3 times with PBS and mounted using Dako Omnis Fluorescence Mounting Medium (Agilent), dried overnight at room temperature and stored at 4 °C. The stained samples were imaged using Olympus CSU-W1 SoRa Spinning Disk Microscope (Olympus) using 20x 0.7 NA air, 40x 1.25 NA and 60x 1.3 NA silicone immersion objectives or STELLARIS 8 Confocal Microscope Platform (Leica Microsystems) using 40x 1.3 NA oil immersion objective.

#### Flow cytometry

For characterization of cell surface markers, cells (2x10^5^ cells per marker) were washed once with PBS and detached using TrypLE. The cells were collected by centrifugation, washed with PBS and stained with fluorochrome-conjugated antibodies in 1% BSA PBS on ice for 1 hour. The cells were washed with PBS, resuspended in 1% BSA PBS and stored on ice prior to analysis. For intracellular staining, cells were fixed with 4% PFA in PBS for 15 min, washed with PBS and permeabilized in Intracellular Staining Permeabilization Wash Buffer (BioLegend) for 1 hour. The cells were collected by centrifugation, washed with PBS, and stained with primary antibody on ice for 1 hour. Cells were washed with PBS and stained with fluorochrome-conjugated secondary antibodies in 1% BSA PBS on ice for 1 hour. The cells were washed with PBS, resuspended in 1% BSA PBS and stored on ice prior to analysis. Cells were analyzed using a BD LSRFortessa™ Cell Analyzer (BD Biosciences) and analyzed using FlowJo software (FlowJo, LLC, v10.8.1). At least 10,000 events per condition were recorded.

#### iLungPro enrichment by fluorescence-activated cell sorting

iLungPro were washed with PBS and detached using TrypLE at 37 °C for 10 minutes. DMEM (Thermo Fisher Scientific) + 10%FBS (Thermo Fisher Scientific) was added to the dissociated cells inactivate to TrypLE. Cells were collected by centrifugation at 300 x g for 5 minutes and resuspended to achieve 10^7^ cells/ml and stained with anti-CPM antibody (FUJIFILM Wako Shibayagi, 1:200) in 10% FBS PBS on ice for 1 hour. Cells were then washed with PBS and stained with Cy3-conjugated secondary antibodies (Thermo Fisher Scientific) in 10% FBS PBS on ice for 1 hour. The cells were washed with PBS, resuspended in sorting buffer (10% FBS PBS; 10 µM Calcein blue, Life Technologies, 10 µM Y- 27632) and stored on ice prior to cell sorting. The cells were sorted using a BD FACSAria™ Fusion Flow Cytometer (BD Biosciences) to enrich CPM+ iLungPro cells into collection buffer (10% FBS PBS; 10 µM Y-27632) kept at 4°C. The sorting quality is checked prior to downstream processes.

#### iVEC enrichment by magnetic cell sorting

iVEC were washed with PBS and detached using Accutase (STEMCELL Technologies) at 37 °C for 5 minutes. DMEM/F12 (Thermo Fisher Scientific) was added to the dissociated cells to dilute out Accutase, cells were collected by centrifugation at 300 x g for 5 minutes and resuspended to achieve 1.2x10^7^ cells/ml and stained with anti-CD144 microbeads in autoMACS^®^ Rinsing Solution (Miltenyi Biosciences) on ice for 20 minutes. The cells were washed with autoMACS^®^ Rinsing Solution and collected by centrifugation at 300 x g for 5 minutes and resuspended in 500 µl autoMACS^®^ Rinsing Solution. The resuspended cells were passed through LS Column (Miltenyi Biosciences) and the retained cells were eluted in autoMACS^®^ Rinsing Solution + 10 µM Y-27632 for downstream culture.

#### RT-qPCR

After the cells were washed once with PBS and lysed in 350 µl RLT buffer (QIAGEN), the total RNA was extracted using RNeasy Mini Kit (QIAGEN). Total RNA was pooled from duplicate wells per condition and stored at -80 °C until further use. The total RNA was measured using a NanoDrop™ One/OneC Microvolume UV-Vis Spectrophotometer (Thermo Fisher Scientific) to assess quality and yield. Conversion to cDNA was achieved using a QuantiTect Reverse Transcription Kit (QIAGEN). RT-qPCR analysis was performed using a QuantStudio 12K Flex Real-Time PCR System (Thermo Fisher Scientific) and TaqMan™ probe-based detection (Thermo Fisher Scientific) in triplicate. The expression of target genes was normalized to house-keeping gene β-actin and fold expression was calculated using ΔΔCt method in comparison to enriched iLungPro cells or a primary Adult Lung sample.

#### LDL uptake assay

iVEC were incubated with 2.5 μg/ml Alexa Fluor™ 594 AcLDL (Thermo Fisher Scientific) in StemPro34 medium for 4 hours at 37°C. Thereafter, cells were washed three times with PBS, detached with TrypLE and fixed with 4% PFA in PBS for 10 minutes at room temperature. Cells were analyzed using a BD LSRFortessa™ Cell Analyzer and analyzed using FlowJo software. At least 10,000 events per condition were recorded.

#### Angiogenesis assay

Angiogenic potential was tested as described before ^50^. Briefly, tissue culture plate was coated with undiluted Matrigel at 150 µl/cm^2^, and iVEC were seeded at 3x10^4^ cells/cm^2^. The formation of tubular structure is inspected 6 hours post iVEC seeding.

#### TER measurements

Trans-barrier electrical resistance (TER) measurements were used to assess the barrier integrity of epithelial and endothelial cells seeded on AX12. The TER measurements were measured using an Epithelial Volt/Ohm Meter (EVOM2 and EVOM3; World Precision Instruments) using a 96-well plate electrode (STX100 Electrode Corning 96; World Precision Instruments) with the connector (EVOM3 legacy probe kit, World Precision Instruments) for EVOM3. 24 hours post epithelial seeding, TER measurements were taken every day until the initiation of the ALI where measurements were then taken every other day. To take the measurements, the electrode was sterilized with 70% v/v ethanol (Thermo Fisher Scientific) and connected to the EVOM2 or EVOM3. After securely placing the AX12 device into the ^AX^Dock, the “TEER Measurement” function was selected on the ^AX^Exchanger. The electrode was positioned between the outlet and central wells and the resistance value was recorded. To measure the background TER, an empty AX12 plate without cells was used. The background value was subtracted from the measured value of a sample and then multiplied by the surface area of the well (0.071cm²) to obtain the final TER reading in Ω·cm^2^.

#### Scanning electron microscopy

Samples were fixed with a mixture of 4% formaldehyde, 1.25 % glutaraldehyde (Merck), 0.04 M Sucrose in 200 mM HEPES (Merck) pH 7.4 overnight at 4°C. Samples were processed using a Biowave Pro (Pelco, USA) as follows: samples were washed twice in 200 mM HEPES pH 7.4 at 250 W for 40 s, post-fixed/contrasted using a solution of 2% osmium tetroxide (Taab) and 1.5% potassium ferricyanide (Taab). Samples were washed with ddH_2_O, and incubated with 1% thiocarbohydrazide (Merck) in distilled water (v/v) for 14 min at 100 W power (with/without vacuum 20 ”Hg at 2 minutes intervals). Samples were then incubated with 2 % osmium tetroxide distilled water (w/v) for 14 minutes at 100 W power (with/without vacuum 20 ”Hg at 2 min intervals) and washed. Samples were incubated in 1 % aqueous uranyl acetate (Agar scientific) in distilled water (w/v) for 14 minutes at 100 W power (with/without vacuum 20 ”Hg at 2 min intervals) and then washed. Samples were dehydrated using a stepwise ethanol series of 50, 75, 90 and 100 % v/v at 250 W for 40 s per step. CPD was performed using an EM CPD300 (Leica Microsystems) and ethanol as the solvent. Samples were coated with 2 nm gold before imaging on a Quanta SEM (FEI, United States).

#### Transmission electron microscopy

Samples were processed using a Biowave Pro (Pelco, USA) with use of microwave energy and vacuum. In the first instance, samples were fixed by adding a double strength mixture of 8% formaldehyde, 2.5 % glutaraldehyde, 0.04 M Sucrose in 200 mM HEPES pH 7.4 v/v to the apical side of the membrane. This was followed by a media exchange to replace the basal medium with mixture of 4% formaldehyde, 1.25 % glutaraldehyde, 0.04 M Sucrose in 200 mM HEPES pH 7.4. Samples were incubated at RT for 5 min before being transferred to fresh single strength fixative and allowed a further 30 minutes at RT before transfer and storage at 4 degrees. Samples were washed twice in 100 mM HEPES at 250 W for 40 s, postfixed using a solution of 2% osmium tetroxide, 1.5% potassium ferricyanide. Samples were washed with ddH_2_O and treated with 1% thiocarbohydrazide in distilled water (v/v) for 14 minutes at 100 W power (with/without vacuum 20 ”Hg at 2 min intervals). Samples were incubated with 2 % osmium tetroxide distilled water (w/v) for 14 minutes at 100 W power (with/without vacuum 20 ”Hg at 2 minutes intervals) and then washed. Samples were incubated in 1 % aqueous uranyl acetate in distilled water (w/v) for 14 minutes at 100 W power (with/without vacuum 20 ”Hg at 2 min intervals). And then incubated with Reynolds’ lead aspartate for 14 minutes at 100 W power (with/without vacuum 20 ”Hg at 2 min intervals). Samples were infiltrated with a dilution series of 25, 50, 75, and 100% Durcupan ACM (Merck) (v/v) resin to ethanol. Each step was 3 min at 250 W power (with/without vacuum 20 Hg at 30-second intervals). Sample were cured for a minimum of 48 h at 60°C. Sample block was orientated for trimming the cells transversely against the membrane. Using a razor blade, excess resin and plastic was removed, then fine trimming is done using a 35° ultrasonic, oscillating diamond knife (DiATOME, Switzerland) set at a cutting speed of 0.6 mm/s. The frequency is set by automatic mode and a voltage of 6.0 V is applied. The knife is installed in a ultramicrotome EM UC7 (Leica Microsystems). 55-70 µm sections were transferred to TEM slot grids with a plioform support film and allowed to dry before imaging by TEM imaging using a JEOL1400FLASH/TEM. Micrographs were visualized and linear adjustments made to contrast and brightness using FIJI (version:2.9.0/1.54f).

#### Image-based single-cell analysis

For the quantification of SP-C, PDPN and ABCA3, the first step in this analysis was to automatically segment the full cytoplasmic extent of each cell. This was achieved by using a ZO-1 membrane marker as an input to the generalist segmentation algorithm ***Cellpose*** ^51^. A comprehensive, whole-cell measurement of fluorescent marker intensity was needed as the expression of SP-C/PDPN/ABCA3 was variable across the cell height in the z-axis. To achieve this, the ***btrack*** ^52^ tracking algorithm was co-opted to link each single-cell instance in each z-slice over the full image volume. During the initial localization step of the ***btrack*** method, the underlying single-cell mean intensities of the cell-type fluorescent markers were recorded for each segment. For manual verification of this analysis, a bespoke ***napari*** ^53^ key binding script was created that allowed for the unbiased selection of a sub-sample of user-chosen positive and negative single-cell examples of each data set. This resulted in a distribution of pixel values for both positively- and negatively expressing cells, resulting in a reliable quantification of single-cell identity on a population level that helped inform the veracity of this approach.

For the quantification of AGER, the image stacks were loaded into FIJI (version:2.9.0/1.54f) using Bio-Formats Importer, Z-projections were generated using maximum projection. An individual nuclei region of interest (ROI) was segmented by thresholding the DAPI channel and quantified as the total cell number. The AGER channel was thresholded to remove background signal, and cells with positive AGER signal on cell membrane contacting all neighboring cells were manually quantified.

#### Image-based single nuclei analysis

The image stacks were loaded into FIJI (version:2.9.0/1.54f) using Bio-Formats Importer, Z-projections were generated using maximum projection. An individual nuclei ROI was segmented by thresholding the DAPI channel and converting the binary image mask to selection. NKX2-1 channel and NKX2-1 fluorescence intensity were measured using Analyze Particles, redirecting the analysis of the ROIs to the other image channels.

#### Library preparation for single-cell RNA sequencing

To dissociate the apical cells, 70 µl of TrypLE was added to the apical chamber of AX12 and 200 µl of TrypLE is flowed into the basal chamber using “medium change” function of ^AX^Exchanger, AX12 was then incubated at 37 °C for 10 min. The apical side was washed once with PBS and then dissociated to single cells by pipette up and down and transferred to 1%BSA PBS. AX12 is then opened and flipped upside down, basal monolayer was dissociated to single cells by pipette up and down and transferred to 1%BSA PBS, the basal side was washed once with PBS and collected. The harvested single cells were collected by centrifugation at 300 x g for 5 mins at 4 °C, washed once in PBS and resuspended in 0.04% BSA PBS, passed through cell strainer (Bel-Art), counted using acridine orange (AO) and propidium iodide (PI) and the Luna-FX7 Automatic Cell Counter (Logos Biosystems). Approximately 20,000 cells were loaded on Chromium Chip and partitioned in nanolitre scale droplets using the Chromium Controller and Chromium Next GEM Single Cell Reagents (10x Genomics, CG000315 Chromium Single Cell 3’ Reagent Kits User Guide (v3.1 - Dual Index)). Within each droplet the cells were lysed, and the RNA was reverse transcribed. The resulting cDNA within a droplet shared the same cell barcode. Illumina compatible libraries were generated from the cDNA using Chromium Next GEM Single Cell library reagents in accordance with the manufacturer’s instructions (10x Genomics, CG000315 Chromium Single Cell 3’ Reagent Kits User Guide (v3.1 - Dual Index)). Final libraries are QC’d using the Agilent TapeStation (Agilent Technologies) and sequenced using the Illumina NovaSeq 6000 Sequencing System (Illumina) using Sequencing read configuration: 28-10-10-90.

#### Bioinformatic analysis of single-cell RNA sequencing

Gene expression was quantified using Cell Ranger count (v.6.0.1) against the prebuilt reference GRCh38-2020-A (10x Genomics). All subsequent analyses were carried out using Seurat (v4) package^54,55^, with default parameters unless specified, in R-4.2.0 (R Core Team (2022) https://www.R-project.org/). Each individual dataset (iAM breathing, iAM static, iLoC breathing, iLoC static) was normalised with the “LogNormalize” method. The top 2,000 highly variable genes were identified using the “FindVariableFeatures” function and the data was scaled (“ScaleData” function). We run PCA selecting the first 20 PCs, constructed a shared nearest neighbour graph using the “FindNeighbors” function and clustered the cells using Louvain clustering (“FindClusters” function) at a resolution of 1.2. Clusters with high average number of mitochondrial content (> 5 percent.mt), low average number of genes (< 1,500 nFeature_RNA) and low number of molecules (< 5,000 nCount_RNA) were removed, and individual samples were reprocessed as before, recalculating the highly variable features, and selecting the first 20 PCs to construct UMAP plots. All four individual datasets were then integrated (“IntegrateData” function) using the Seurat standard CCA integration workflow, after identifying 2,000 anchor features. The resulting integrated data was visualized on the UMAP space using the first 50 PCs.

Integrated clusters at resolution 1 were annotated based on a combination of the expression of known marker genes for each cell type, and top cluster specific gene markers identified using the “FindAllMarkers” function (with the settings: only.pos = TRUE, logfc.threshold = 0.25, min.pct = 0.2). The integrated dataset was also compared to a reference Human Lung Cell Atlas (HLCA) of the healthy respiratory system spanning over 584,000 cells of 61 cell identities ^43^. To assess similarities between them, we used 1) the cell label transfer method and projection of query cells onto reference UMAP structure (“FindTransferAnchors” and “MapQuery” functions) from Seurat, and 2) the cluster label assignment from “clustifyr” (v1.10.0) that adopts a Spearman correlation-based method to find reference transcriptomes with the highest similarity to query cluster expression profiles ^56^. The iPSC-derived macrophage (iPSDM) population identified in the integrated data was also sub-setted for further investigation. iPSDM cells were extracted from the individual samples, reprocessed, and integrated again as described above and subclustered at resolution 0.6. Differential expression analysis was performed using the “FindMarkers” function (DESeq2 test without logfc or minimum percentage expressed thresholds). The Seurat DESeq2DETest function was modified to output the Wald statistic. Gene Set Enrichment Analysis (GSEA) ^57^ was performed using “ClusterProfiler” v.4.6.2 ^58^. Gene lists from the differential expression analysis for each comparison were ranked by the Wald statistic. C2- CP:REACTOME from the MSigDB collection (MSigDB v2023.1) was assessed with the parameters minGSSize = 5 and maxGSSize = 5000, and only pathways with an adjusted p-value lower than 0.05 were considered statistically significant^59^. Dot plots illustrating selected pathways were made with ggplot2.

#### Cytokine measurements

Apical and basal media of AX12 were harvest and centrifuged through 0.22 µm filter column (Thermo Fisher Scientific) twice to remove cells and Mtb. The media were stored at -80 °C prior to cytokine analysis. Cytokine screening was performed using Bio-Plex Pro Human Cytokine Screening Panel, 48- Plex (BioRad) on a Bio-Plex^®^ 200 System (BioRad) following the manufacturer’s protocol.

#### RNAScope imaging and analysis

Sample were fixed using 4% buffered PFA. AX12 membranes were recovered using a heated scalpel to remove excess plastic and equilibrated in an increasing sucrose gradient of 10%, 20% and 30% in PBS at 4°C. Membranes were then embedded in OCT medium (Agar scientific) with dry ice. Cryosections were cut to a thickness of 12 µm at an OT of -18 °C and a CT of -19 °C using a CM3050S Cryostat (Leica microsystems). Slides were stored at -80 °C until use. The 12 µm fixed frozen sections were stained on the BOND RX Fully Automated Research Stainer (Leica microsystems) using RNAscope™ LS Multiplex Fluorescent assay (Bio-Techne) applying a 10 min protease treatment at room temperature and using target probes *IL-6* (Bio-Techne), *IL-8* (Bio-Techne) with Opal 570 and Opal 780 TSA fluorophores (Akoya Biosciences). Snapshots of each membrane were taken using the Vectra Polaris™ Automated Quantitative Pathology Imaging System (Akoya Biosciences). Subsequent analysis involved the use of QuPath Software Version 0.4.4. to perform perinuclear segmentation via DAPI staining. After a manually defined threshold was chosen for each mRNA probe. The QuPath ‘detection measurements’ function was used to segment cellular puncta. The number of positive dots per cell for each probe was recorded.

#### Mtb infection

Mtb H37Rv cryovials were thawed and cultured to late log phase in Middlebrook 7H9 broth medium (BD Biosciences) with 50 µg/ml Hygromycin B (Thermo Fisher Scientific). The Mtb culture is collected by centrifugation at 3000 x g for 5 minutes, washed twice with PBS and collected again by centrifugation. The bacterial pellet is dissociated by vortexing with glass beads for 1 min. The dissociated bacteria were resuspended in Alveolar medium and centrifuged at 1200 x g for 5 min. The supernatant containing single bacterium are collected and measured using a spectrophotometry at OD_600_. The Mtb is diluted in Alveolar medium and infected the iLoC from apical side aiming to achieve MOI = 0.01 without washout until experimental endpoint. After infection, apical and basal media were harvested and the iLoC were fixed as designated timepoints with 4% PFA in PBS at 4 °C overnight.

#### Quantification of Mtb in iLoC

For the quantification of Mtb in iAE and iPSDM at infection timepoints of 2 h and 48 h pi (i.e. intact ZO-1 barrier), the iAE were automatically segmented using a ZO-1 membrane marker as input to ***Cellpose*** ^51^ and the iPSDM were manually segmented using ***napari*** ^53^. The Mtb channel was set with an intensity threshold to visualize only the bacteria, the Mtb area per segmented cell were then measured. For the quantification of total Mtb in iLoC at infection timepoint of 2 h, 48 h and 120 h pi, the image stacks were loaded into FIJI (version:2.9.0/1.54f) using Bio-Formats Importer, Z-projections were generated using maximum projection. Individual Mtb ROI was segmented by thresholding the Mtb channel, converted to mask and the ROI areas were measured using Analyze Particles.

#### Live/Dead Orange staining of iLoC

Thirty minutes prior to the designated timepoint of iLoC sampling, LIVE/DEAD™ Fixable Orange (602) Viability Kit (Thermo Fisher Scientific) was added to apical and basal media. At designated timepoints, iLoC were harvested as described above. Harvested samples were permeabilized with 0.01% saponin in 1%BSA PBS for 1 hour at room temperature, then blocked in 3% BSA PBS for 1 hour. Cells were washed 3 times with PBS, then stained with DAPI (1:10000) and AlexaFlour488-anti-ZO-1 antibody (Thermo Fisher Scientific, #339188, 1:100) in 1% BSA PBS for 1 hour at room temperature. Cells are then washed 3 times with PBS and mounted using Dako Omnis Fluorescence Mounting Medium, dried overnight at room temperature, and stored at 4 °C. The stained samples were imaged using Olympus CSU-W1 SoRa Spinning Disk Microscope using 20x 0.7 NA air objectives, and LIVE/DEAD-positive cells were manually quantified.

#### Quantification and statistical analysis

All data presented in this work were obtained from at least n = 2 independent experiments. Statistical analyses applied are described in each figure legend. For the comparison of two groups, unpaired Student’s t-test was used (Fig. 3g, Supplementary Fig. 4c-d). For the comparison of multiple groups with one variable, one-way Analysis of Variance (ANOVA) with Tukey’s Honestly Significant Difference test was used (Fig. 5e-f, 5h-j, 6b-h). *P* < 0.05 was considered statistically significant.

Statistical significance was determined using Prism v.10.0.1 software (GraphPad).

## Data Availability

Single-cell RNA-seq datasets have been deposited at GEO (GSE252601) and are publicly available as of the date of publication. Accession numbers are listed in the key resources table. Microscopy data reported in this paper will be shared by the lead contact upon request. All original code has been deposited at GitHub and is publicly available as of the date of publication. DOIs are listed in the key resources table. Any additional information required to reanalyze the data reported in this work paper is available from the lead contact upon request.

## Code Availability

All original code has been deposited at GitHub and is publicly available as of the date of publication.

## Acknowledgements

We are indebtedly thankful to Annie Moisan and Cara Buchanan (HOPE, Wellcome LEAP) for her encouragement and support throughout this project. Work on the Lung Engineers project was generously supported by Wellcome Leap funding as part of the HOPE Program. We thank the Human Embryonic Stem Cell Unit, Advanced Light Microscopy, and Advance Sequencing facilities at the Crick for their support in various aspects of the work. This work was supported by the Francis Crick Institute (to M.G.G.), which receives its core funding from Cancer Research UK (CC2081), the UK Medical Research Council (CC2081), and the Wellcome Trust (CC2081). This project has received funding from the European Research Council (ERC) under the European Union’s Horizon 2020 research and innovation programme (grant agreement n° 772022). For the purpose of Open Access, the author has applied a CC BY public copyright license to any Author Accepted Manuscript version arising from this submission. Figures created in BioRender. Luk, C.H. (2024) URL

## Author Contributions

Conceptualization, M.G.G., C.H.L; methodology, C.H.L., G.C., A.F., E.P., N.H., J.D.S; software, I.R.H., N.J.D., R.D.; formal analysis, C.H.L.; investigation, C.H.L., G.C., K.J.G., A.F., N.J.D., N.A.; resources, E.P., J.D.S., NH.; data curation, C.H.L., I.R.H., N.J.D., R.D.; writing – original draft, M.G.G.; writing – review & editing, all authors; visualization, C.H.L., I.R.H., N.J.D.; supervision, M.G.G., C.H.L.; project administration, M.G.G., C.H.L.; funding acquisition, M.G.G.

## Competing interests

N.H. holds minor equity in AlveoliX AG. N.H. is employed by AlveoliX AG.

The remaining authors declare that the research was conducted in the absence of any commercial or financial relationships that could be construed as a potential conflict of interest.

Supplementary Table 1. | Excel file containing differential expression analysis data too large to fit in a PDF.

---

**Supplementary Fig. 1.**
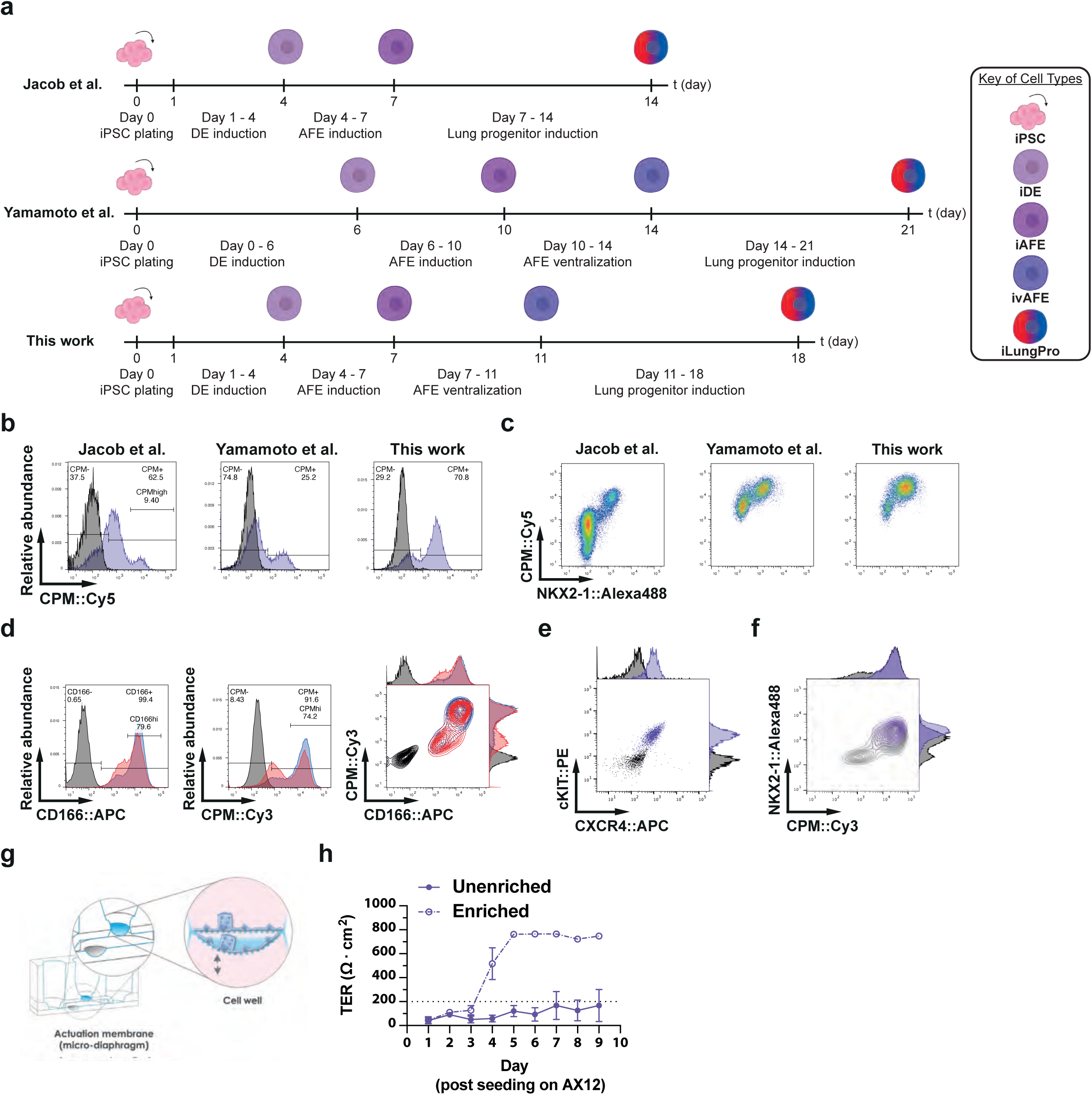
iPSC differentiation into iLungPro and iAT2&iAT1. **a**, Schematic diagram comparing iLungPro differentiation protocols from Jacob et al. 2017, Yamamoto et al. 2017 and this work. **b**, Expression of CPM in iLungPro using different differentiation protocol, signal measured by flow cytometry, representative plot showing isotype (grey) and stained (purple) samples using unit area for y-axis. **c**, Expression of CPM and NKX2-1 in iLungPro using different differentiation protocol, signal measured by flow cytometry; n = 3 independent experiments. **d**, Expression of CD166 and CPM in iLungPro using different cell splitting ratio, signal measured by flow cytometry, representative plot showing isotype (grey), 1:1 split (red) and 1:3 split (blue) samples using unit area for y-axis; n = 3 independent experiments. **e**, Expression of CXCR4 and cKit in iPSC-derived definitive endoderm measured by flow cytometry, representative plot showing isotype (grey) and stained (purple) samples; n = 3 independent experiments. **f**, Expression of CPM and NKX2-1 in iLungPro measured by flow cytometry, representative plot showing unsorted (grey) and CPM+ enriched (purple) samples; n = 3 independent experiments. **g**, Schematic diagram highlighting the 3D mechanical stretching function of AX12. h, TER quantification of iAE on AX12 up to 9 days post-seeding, using unenriched (solid line) or FACS-enriched (dotted line) iLungPro; mean ± s.d., n = 2 independent experiments.

**Supplementary Fig. 2.**
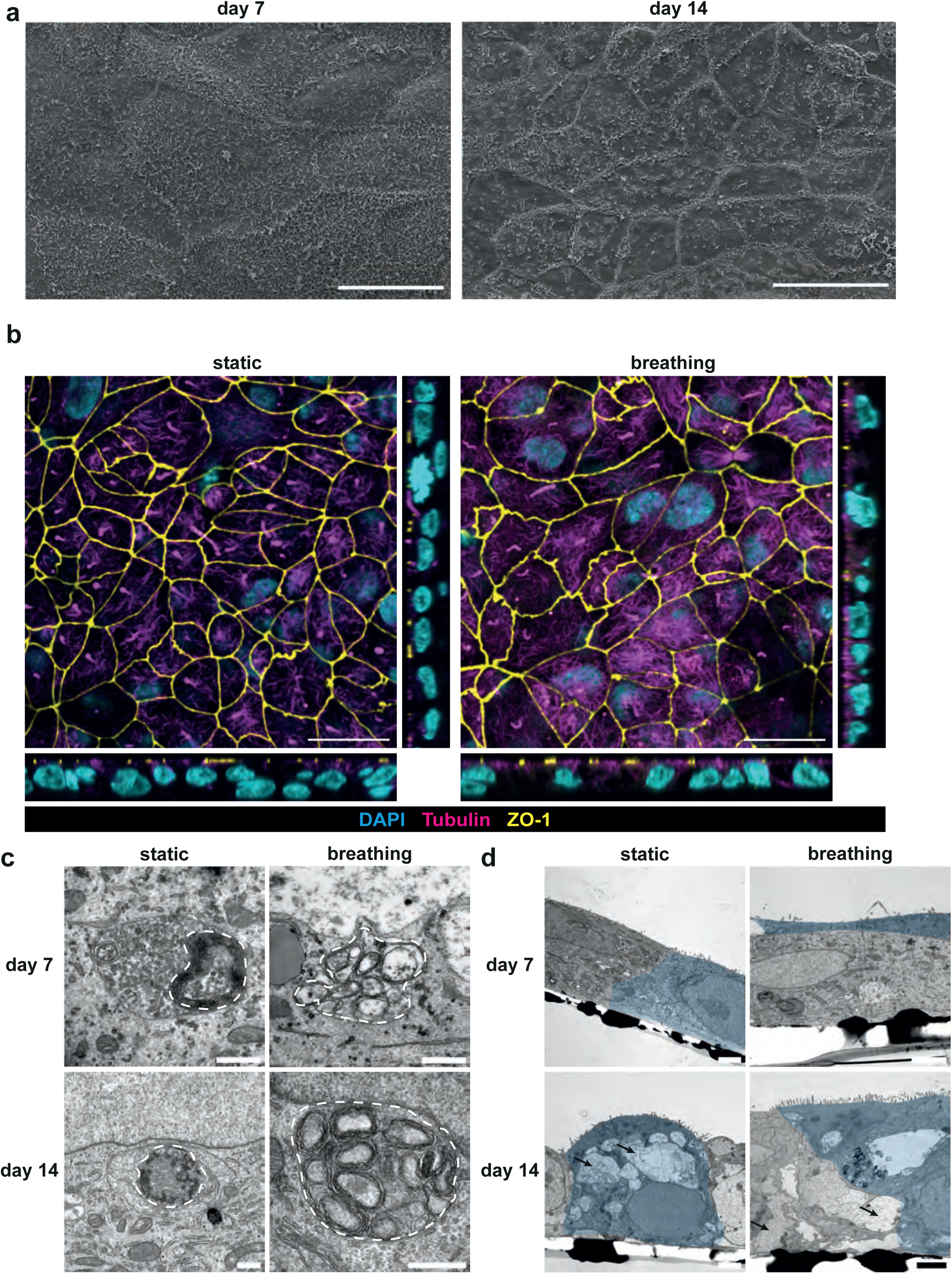
Ultrastructural details of differentiated iAT2 and iAT1s. **a**, SEM image of iAT2 and iAT1 differentiating on AX12 at 7 days post-seeding (left) and 14 days. Scalebar, 20 µm. **b**, Representative confocal images of orthogonal and planar view of the iAE on AX12 showing nuclei (cyan), acetylated Tubulin (magenta) and ZO-1 (yellow) at day 7 post-seeding, under static or breathing conditions. Scalebar, 20 µm. **c**, TEM images of lamellar bodies highlighted by dotted lines in iAT2 at day 7 and day 14 post-seeding, with or without mechanical stretch. Lamella bodies are depicted by dotted lines. Scalebar, 500 nm. **d**, TEM images of ciliated iAT2 and iAT1 cells highlighted in blue at day 7 and day 14 post-seeding, with or without mechanical stretch. Glycogen lake-like structures are indicated by arrows. Scalebar, 2 µm.

**Supplementary Fig. 3.**
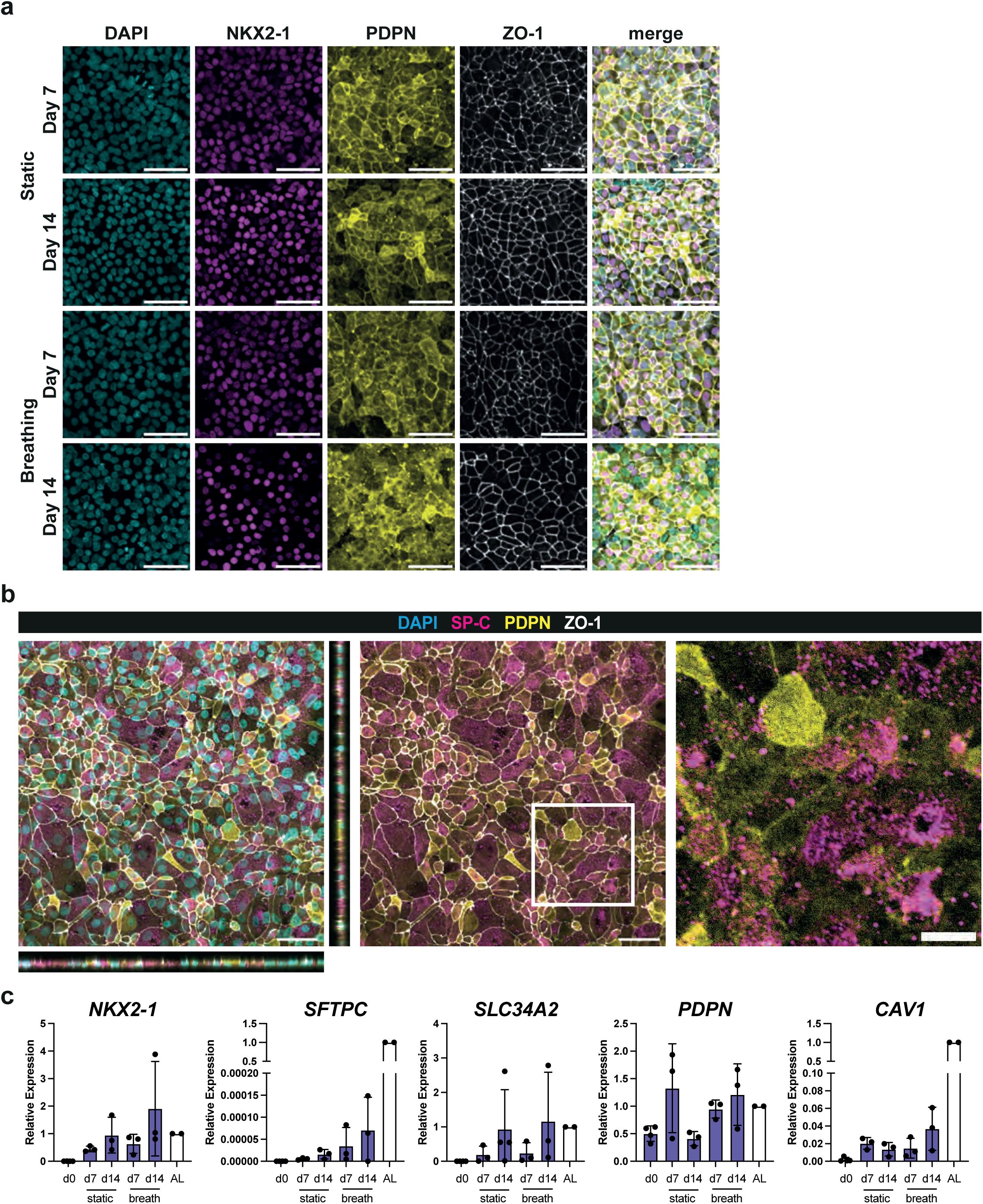
iAT2 and iAT1s in a Lung-on-chip microfluidic device are functional. **a**, Representative confocal images of the iAE on AX12 showing nuclei (cyan), NKX2-1 (magenta), PDPN (yellow) and ZO-1 (white) at day 7 and day 14 post-seeding, under static or breathing conditions. Scalebar, 50 µm. **b**, Representative orthogonal view and magnified view of iAE on AX12 showing nuclei (cyan), mature SP-C (magenta), PDPN (yellow) and ZO-1 tight junctions (white) at day 14 post-seeding, under static conditions. Magnified view depicts square in the middle panel. Scale bar, 50 µm (orthogonal view) and 20 µm (magnified view). **c**, qPCR quantification of marker gene expression levels of iAT2 and iAT1, *NKX2-1*, *SFTPC*, *SLC34A2*, *PDPN* and *CAV1* in lung progenitors (d0), iAE cells at day 7 and day 14 post-seeding, under static or breathing conditions and primary adult lung (AL); mean ± s.d., n = 3 independent experiments.

**Supplementary Fig. 4.**
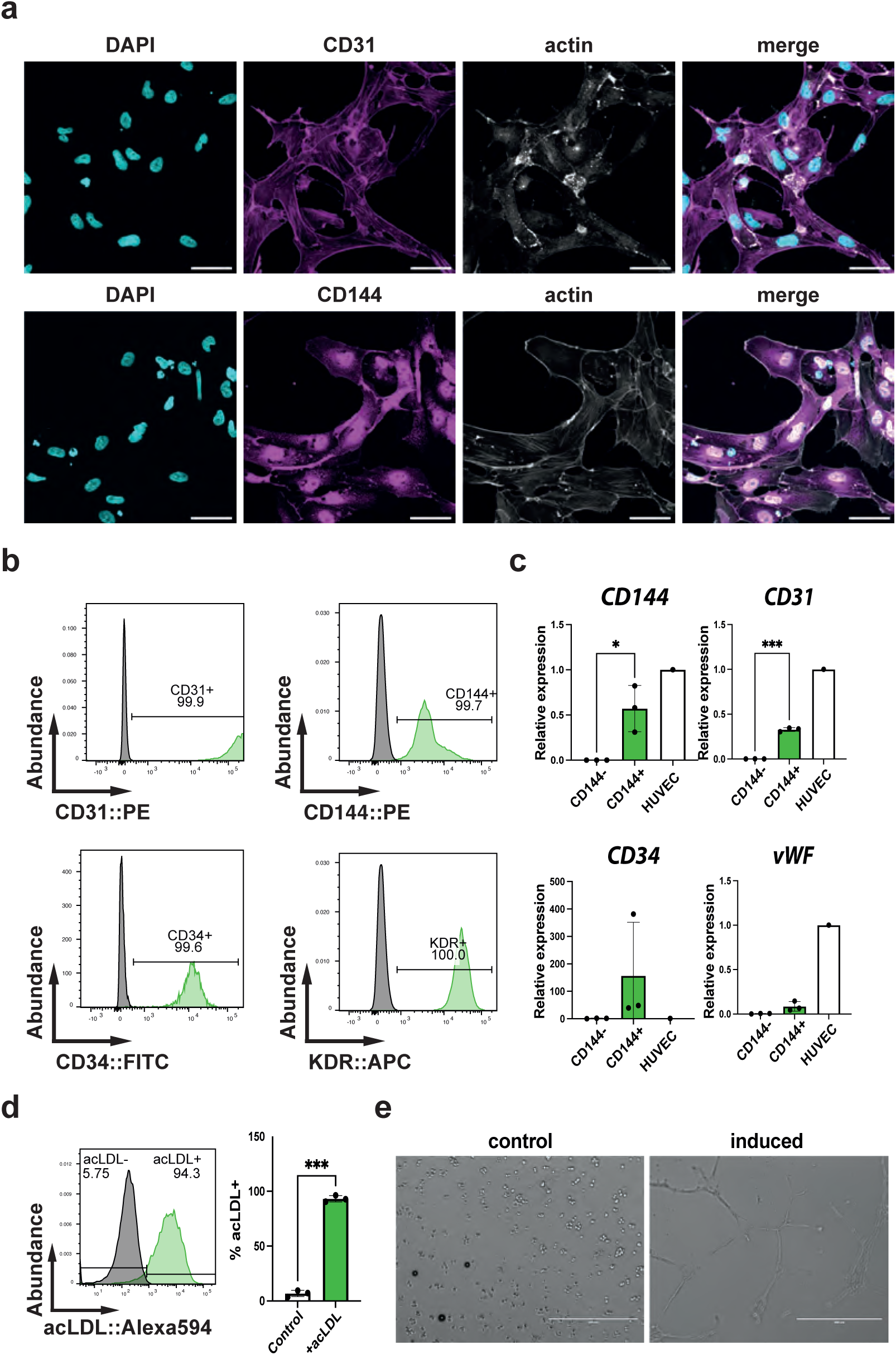
Differentiation and characterization of iVEC. **a**, Representative confocal images of iVEC cultured on glass showing nuclei (cyan), CD31(top, magenta) or CD144 (bottom, magenta) and actin (white). Scalebar, 50 µm. **b**, Expression of CD31, CD144, CD34 and KDR in iVEC measured by flow cytometry, representative plots showing isotype (grey) and stained (green) samples; n = 3 independent experiments. **c**, Quantification of endothelial marker genes expression in iVEC by qPCR, graphs showing flowthrough cells of magnetic sorting (CD144-), magnetic enriched cells (CD144+) and HUVECs; mean ± s.d., n = 3 independent experiments, Student’s t-test). **d**, Quantification of acLDL uptake by iVEC using flow cytometry, representative plots showing control (grey) and acLDL-treated (green) samples. Bar plot showing the quantification of three independent experiments; mean ± s.d., n = 3 independent experiments, Student’s t-test. **e**, Representative brightfield images of iVEC undergoing angiogenesis, showing iVEC in control (Left) and induced (Right) conditions. Scalebar, 400 µm. p-value by unpaired t-test: * P < 0.05, *** P < 0.001.

**Supplementary Fig. 5.**
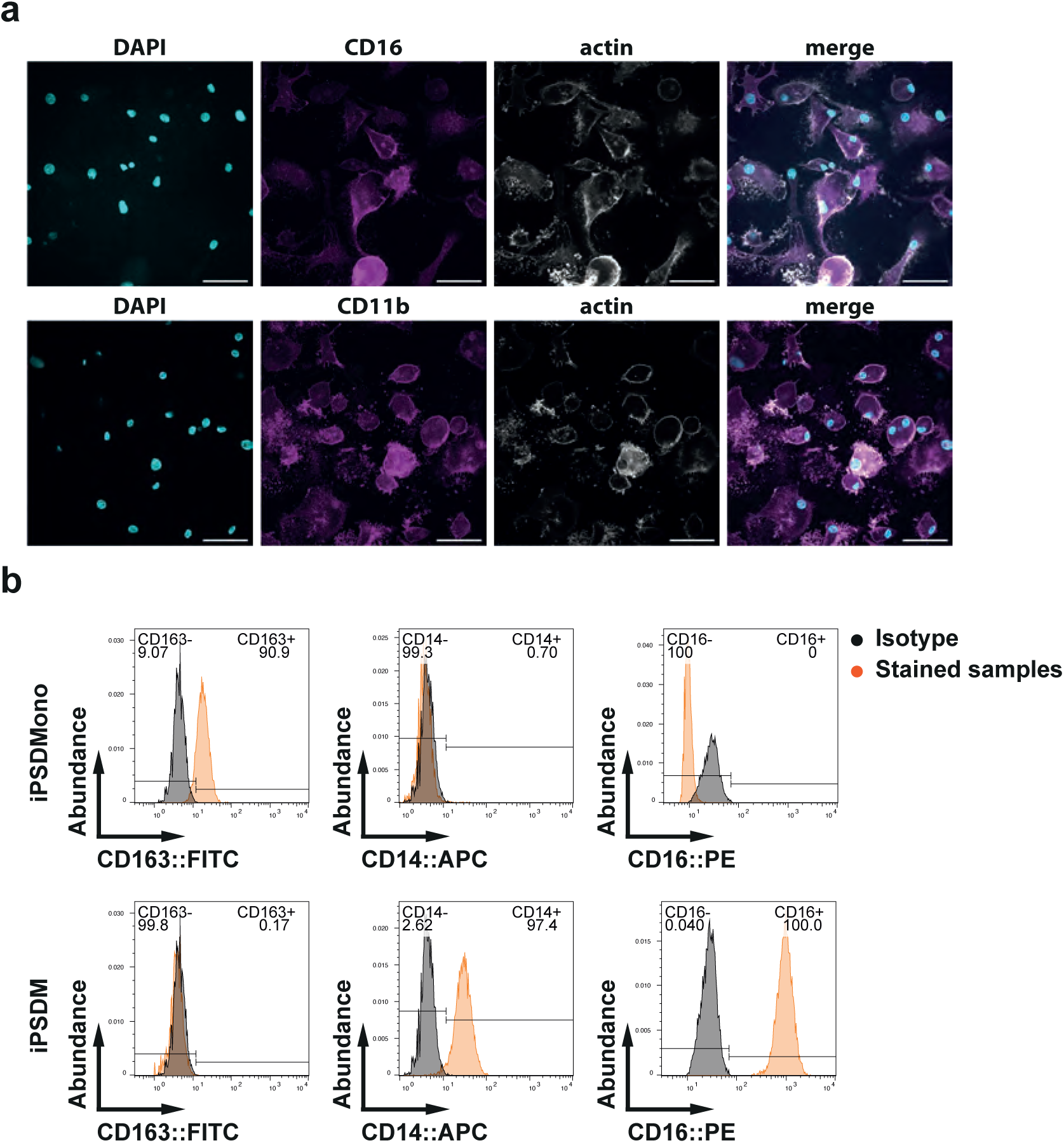
Differentiation and characterization of iPSDM. **a**, Representative confocal images of iPSDM cultured on glass showing nuclei (cyan), CD16 or CD11b (magenta) and actin (white). Scale bar, 50 µm. **b**, Expression of CD163, CD14 and CD16 in iPSDMono (top) and iPSDM (bottom) measured by flow cytometry, representative plots showing isotype (grey) and stained (orange) samples; n = 3 independent experiments.

**Supplementary Fig. 6.**
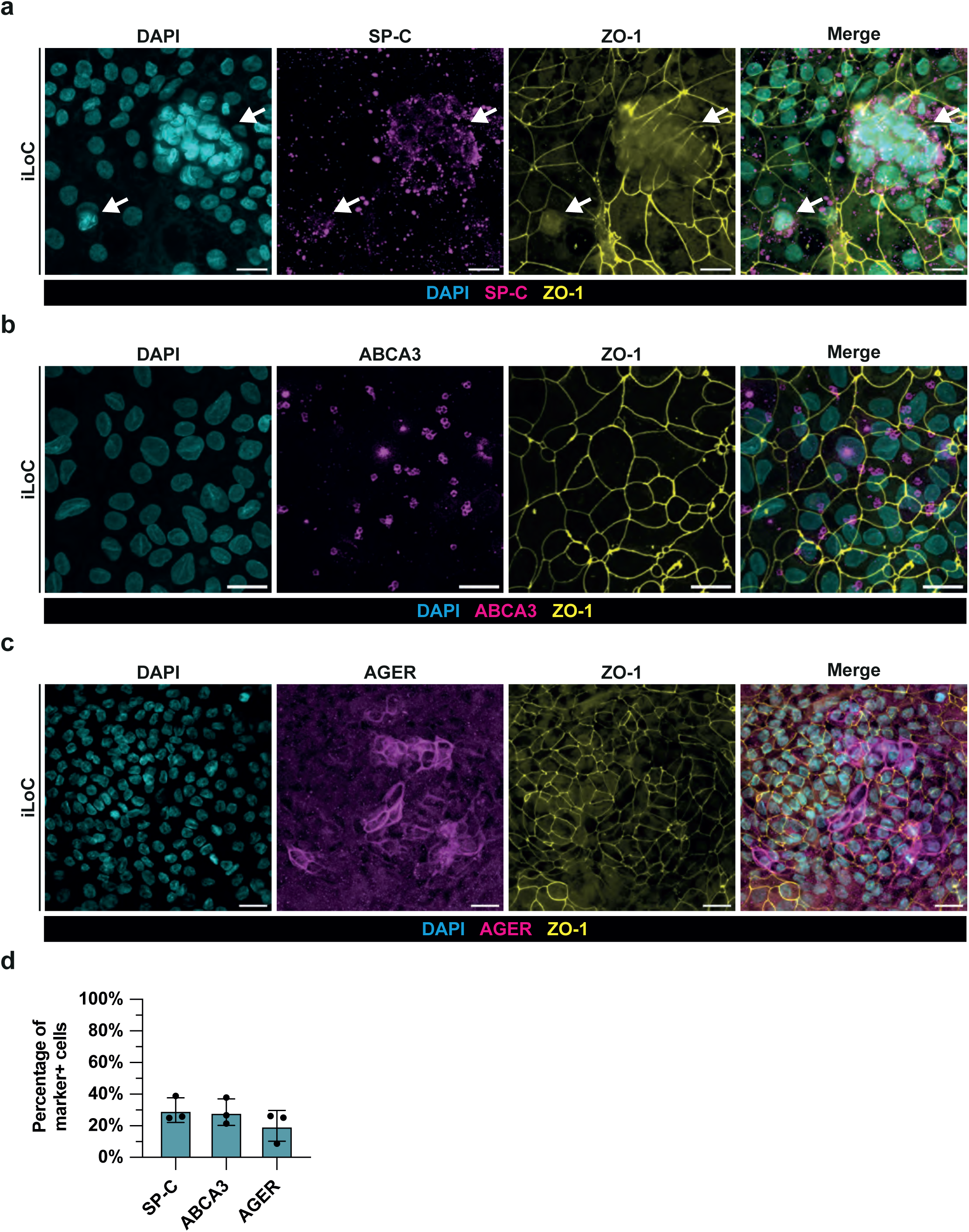
Expression of iAT2 and iAT1 markers of iLoC. **a**, Representative confocal images of iLoC showing nuclei (cyan), SP-C (magenta) and ZO-1 (yellow) under static condition at day 21 of chip assembly, macrophages are depicted by arrows. Scalebar, 20 µm. **b**, Representative confocal images of iLoC showing nuclei (cyan), ABCA3 (magenta) and ZO-1 (yellow) under static condition at day 21 of chip assembly. Scalebar, 20 µm. **c**, Representative confocal images of iLoC showing nuclei (cyan), AGER (magenta) and ZO-1 (yellow) under static condition at day 21 of chip assembly. Scalebar, 20 µm. **d**, Quantification of SP-C, ABCA3 and AGER-expressing cells in under static condition at day 21 of chip assembly; mean ± s.d., n = 3 independent experiments.

**Supplementary Fig. 7.**
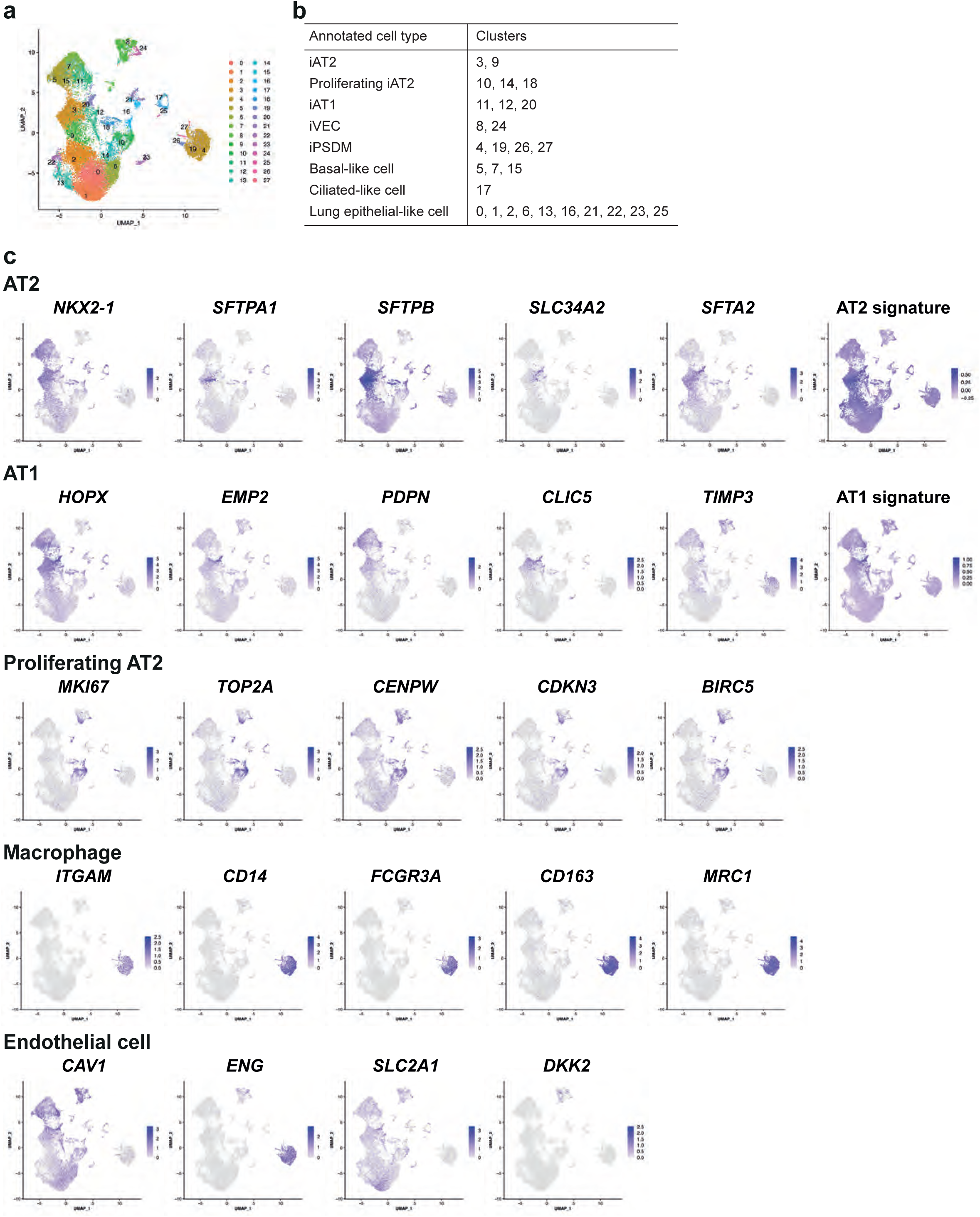
Clustering and cell annotation of iLoC single cell transcriptome analysis. **a**, Uniform Manifold Approximation and Projection (UMAP) showing clusters at resolution 1 for the integrated analysis of 26907 iAM or iLoC-derived cells in the presence or absence of mechanical stretch. (N=1, n=1 per group). **B**, Cell annotation of clusters formed in a,. c, Feature plots showing the expression of AT2 marker genes, *NKX2-1*, *SFTPA1*, *SFTPB*, *SLC34A2*, *SFTA2*, AT2 signature described in Burgess et al. 2024; AT1 marker genes, *HOPX*, *EMP2*, *PDPN*, *CLIC5, TIMP3,* AT1 signature described in Burgess et al. 2024; proliferating AT2 marker genes, *MKI67*, *TOP2A*, *CENPW*, *CDKN3*, *BIRC5*; macrophage marker genes, *ITGAM*, *CD14*, *FCGR3A*, *CD163*, *MRC1*; and endothelial cell marker genes, *CAV1*, *ENG*, *SLC2A1* and *DKK2* as presented in Fig. 4c.

**Supplementary Fig. 8.**
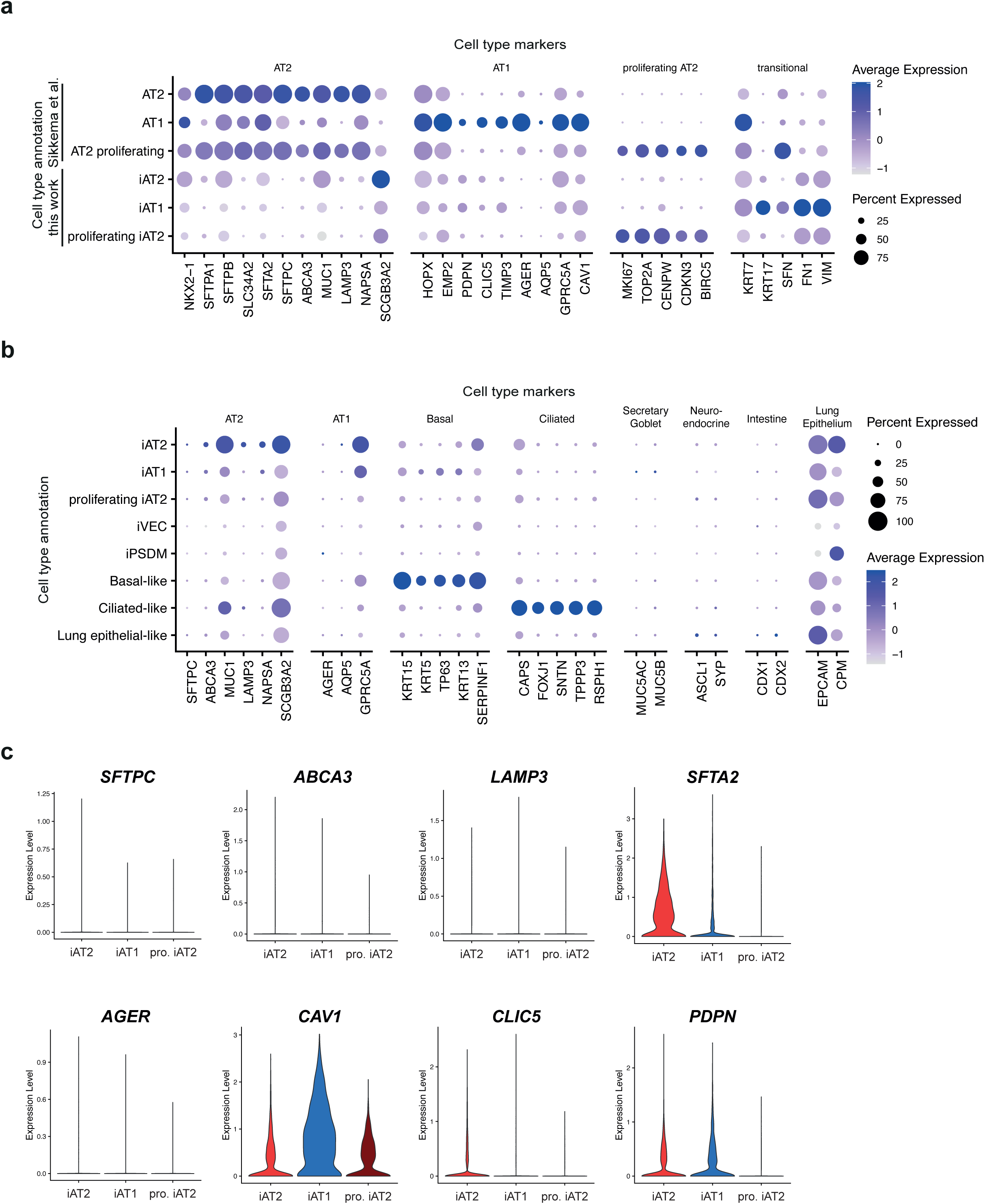
Expression tissue and cell type-specific markers of iLoC. **a**, Dot plot representation of the expression of AT2, AT1, proliferating AT2 and transitional marker genes across AT2, AT1, AT2 proliferating clusters of Sikkema et al. 2023 and annotated iAT2, iAT1 and pro. iAT2 clusters of AX12-derived cells from this study. **b**, Dot plot representation of the expression of additional AT2, AT1, basal cell, ciliated cell, secretary/goblet cell, neuroendocrine, intestinal, and lung epithelial cell gene markers across annotated cell types. **c**, Violin plots of the expression of AT2 marker genes, *SFTPC*, *ABCA3*, *LAMP3*, *SFTA2*; and AT1 marker genes, *AGER*, *CAV1*, *CLIC5*, *PDPN* in annotated iAT2, iAT1 and pro. iAT2 clusters of AX12-derived cells from this study.

**Supplementary Fig. 9.**
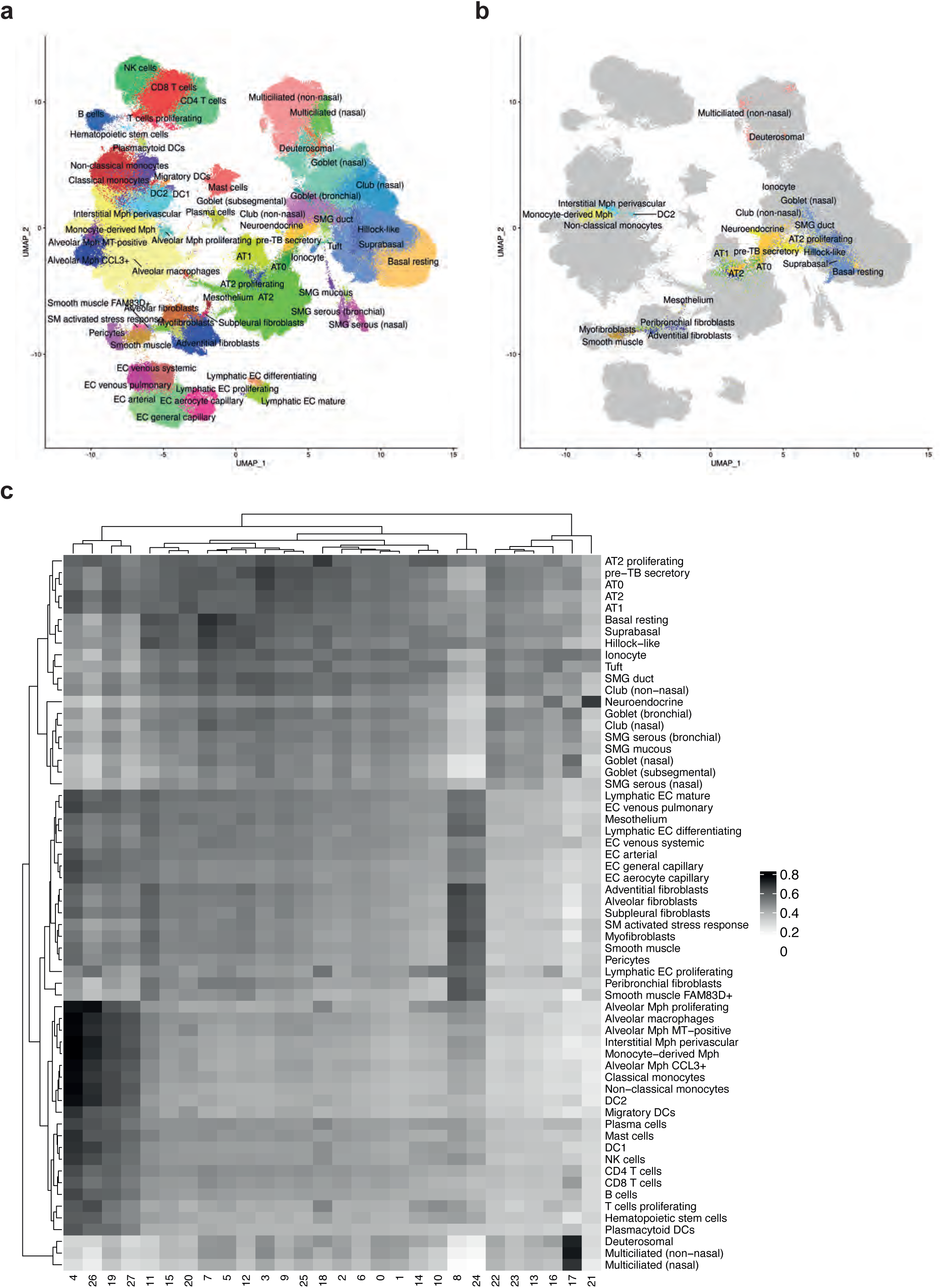
Mapping of iLoC single cell transcriptome to reference dataset. **a**, UMAP visualization of cell identity annotation of the integrated healthy Human Lung Cell Atlas (HLCA) reference dataset published by Sikkema et al. 2023. **b**, UMAP projection of iAM or iLoC-derived cells in the presence or absence of mechanical stretch (query cells) onto the reference HLCA UMAP in **a**. **c**, Heatmap showing spearman correlation coefficients between reference HLCA cell types from Sikkema et al. 2023 and AX12-derived cells (query clusters at resolution 1) from this study.

**Supplementary Fig. 10.**
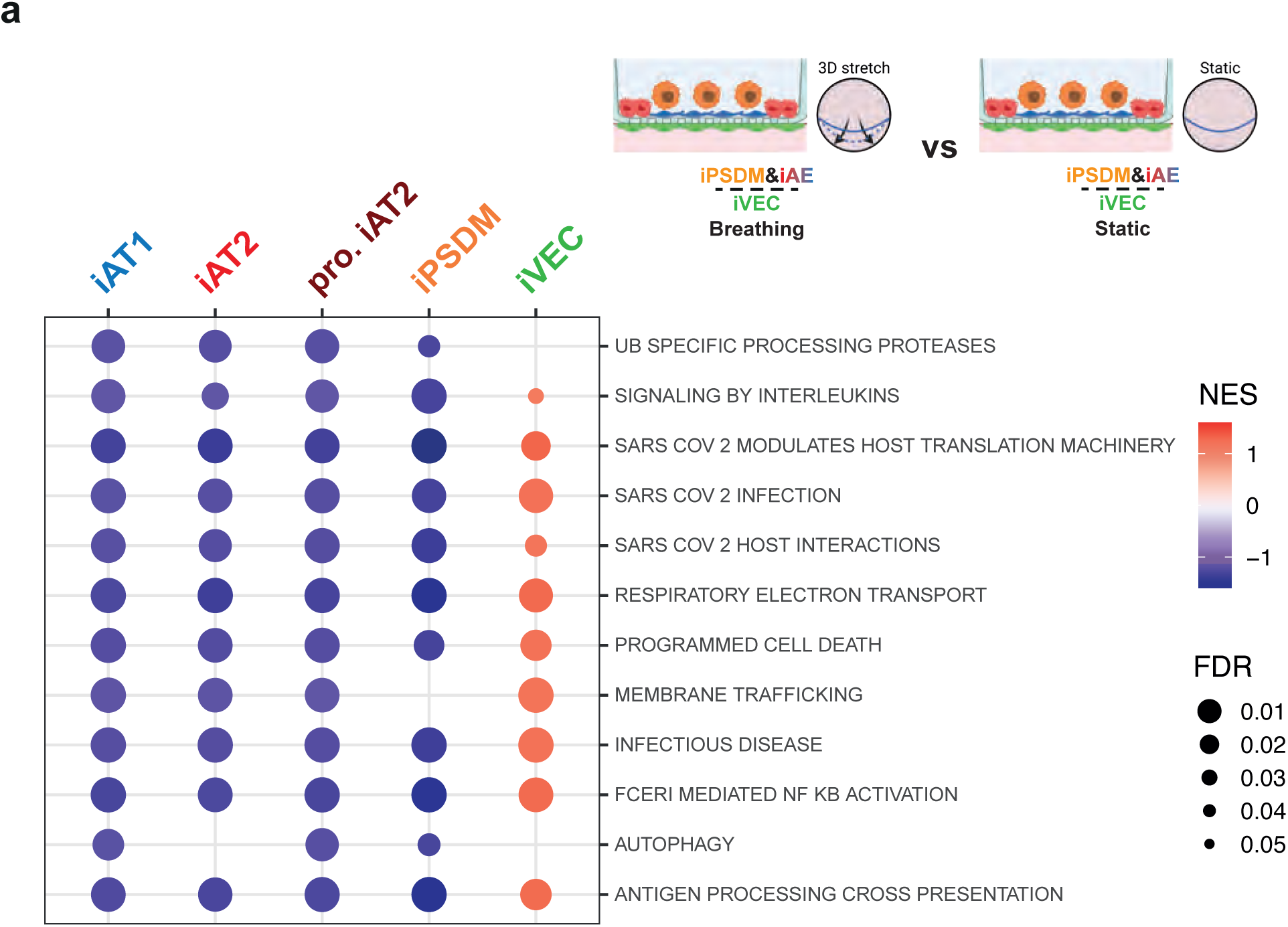
Pathway analysis of iLoC. **a**, Dot plot showing selected Reactome enriched pathways from GSEA analysis for differential expression analysis in iAT2, iAT1, pro. iAT2, iPSDM and iVEC cell types from iLoC under breathing and static condition (Breathing vs Static).

**Supplementary Fig. 11.**
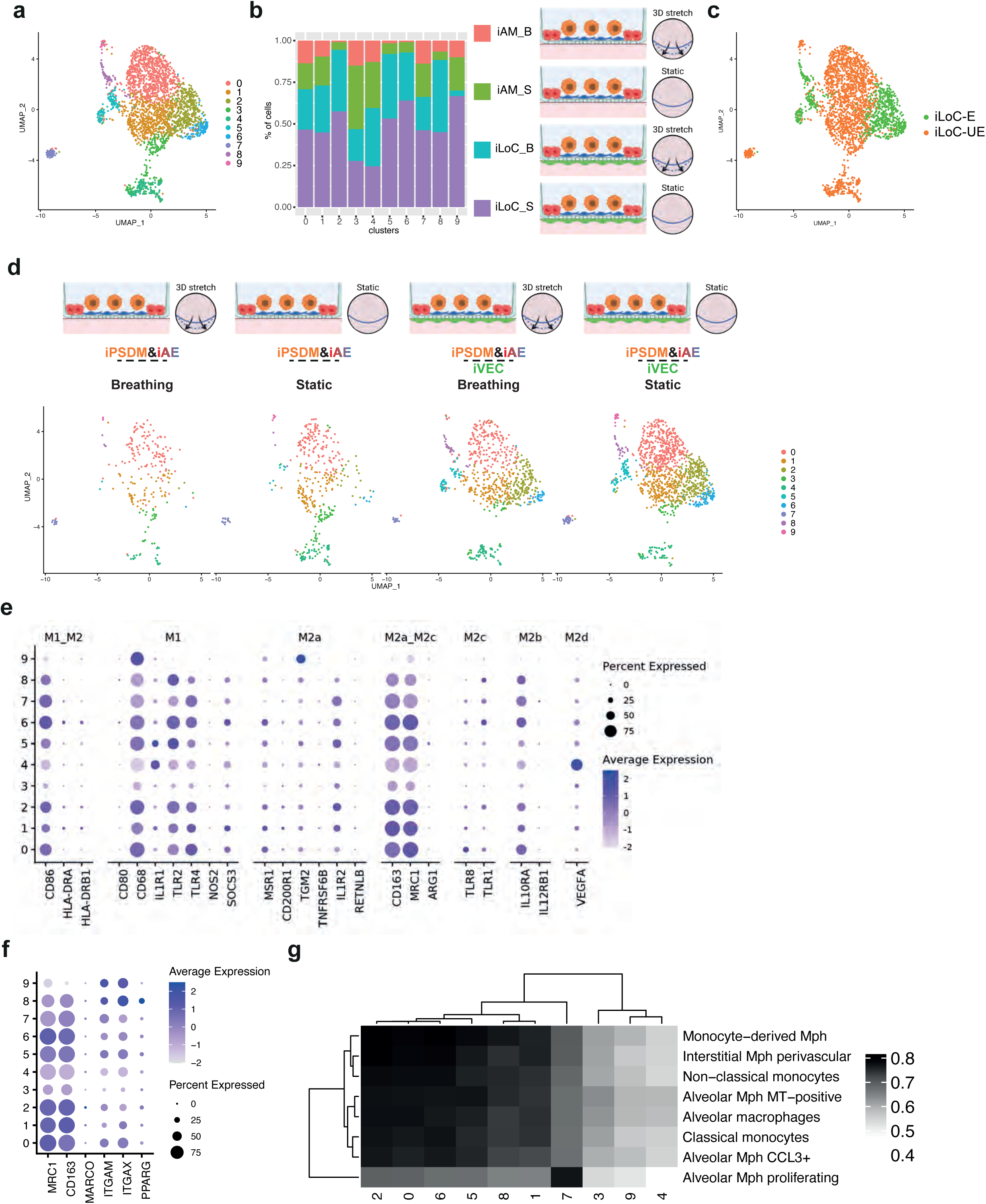
Clustering and subpopulation analysis of iPSDM on iLoC. **a**, UMAP visualization of subclusters identified from iPSDM population from iLoC. **b**, Cell proportions of iPSDM subclusters in each individual sample from **a**. **c**, UMAP visualization of subclusters identified from iPSDM population from iLoC, highlighting iLoC enriched (green) and unenriched (orange) subclusters. **d**, UMAP visualization of iPSDM subclusters from **a** split by experimental condition. **e**, Dot plot representation of the expression of selected macrophage phenotype genes across all iPSDM identified subclusters. **f**, Dot plot representation of the expression of selected alveolar macrophage marker genes across all iPSDM identified subclusters. **g**, Heatmap showing spearman correlation coefficients between macrophage cell types from HLCA from Sikkema et al. 2023 and iPSDM subclusters from this study.

**Supplementary Fig. 12.**
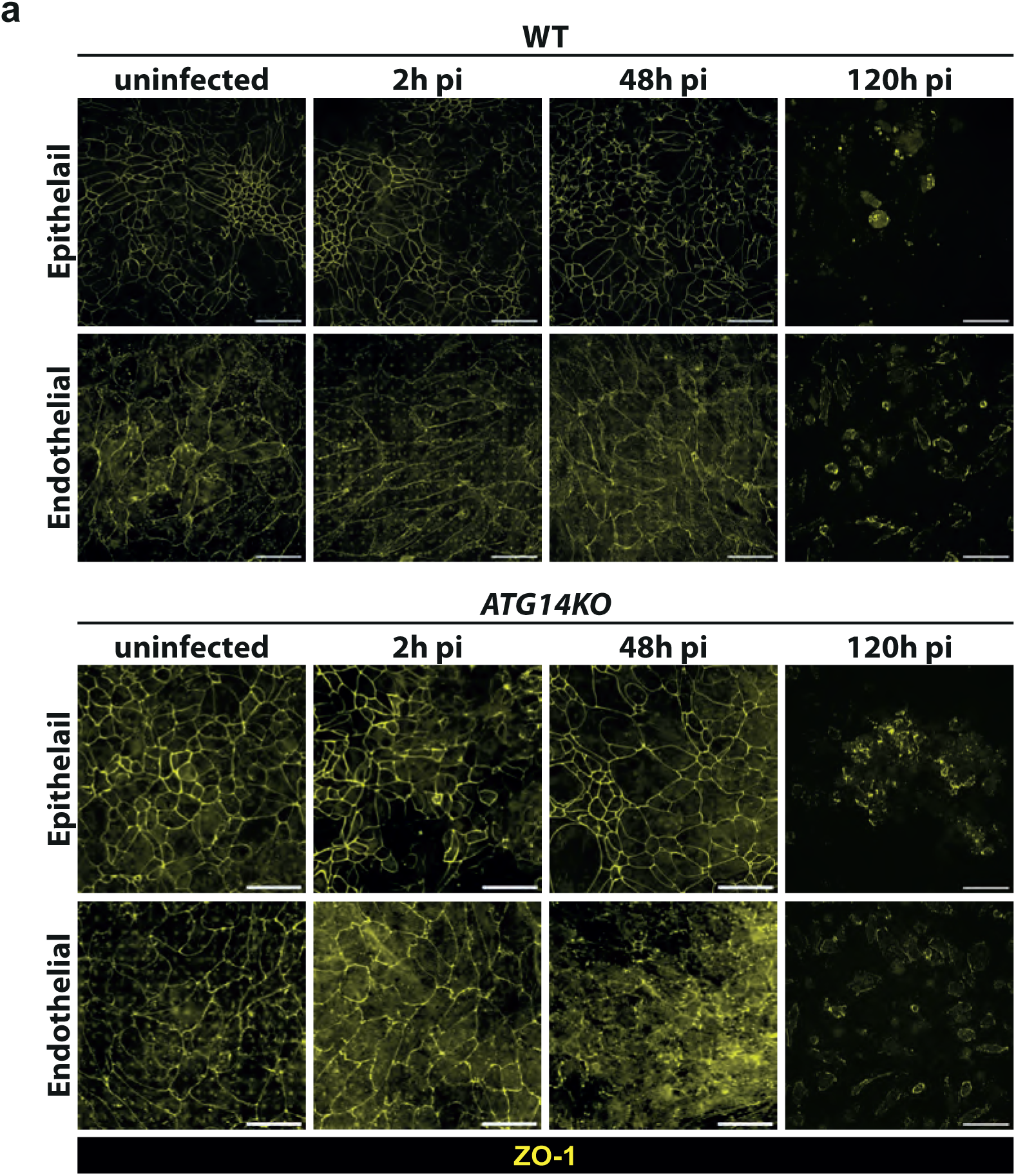
Epithelial and Endothelial barrier integrity in the WT-iLoC and *ATG14KO* GE-iLoC after infection with *M. tuberculosis*. **a**, Representative confocal images of tight junctions of epithelial and endothelial monolayer in iLoC or GE-iLoC marked by ZO-1, images showing ZO-1 (yellow). Scalebar, 50 µm.

**Supplementary Fig. 13.**
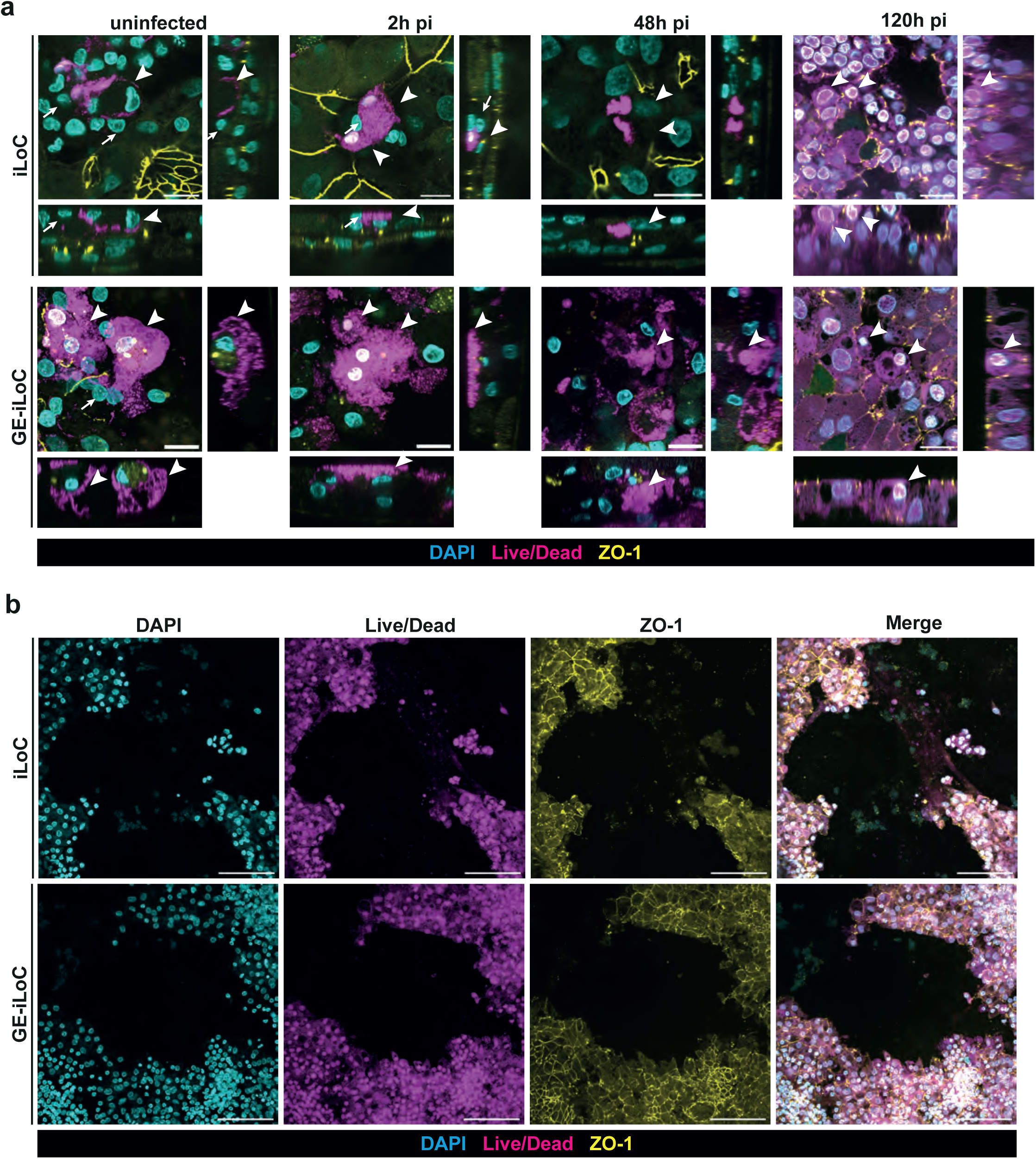
Cell death in the iLoC and *ATG14KO* GE-iLoC after infection with *M. tuberculosis*. **a**, Representative confocal images of cell death in iPSDM in iLoC or GE-iLoC indicated by Live/Dead Orange, images showing nuclei (cyan), Live/Dead Orange (magenta) and ZO-1 (yellow) under uninfected condition, 2 h, 48 h and 120 h pi. Live and dead cells were depicted by arrow and arrowhead, respectively. Scalebar, 20 µm. **b**, Representative images of cell death in iPSDM in iLoC or GE-iLoC indicated by Live/Dead Orange, images showing nuclei (cyan), Live/Dead Orange (magenta) and ZO-1 (yellow) at 120 h pi. Scalebar, 100 µm.

**Supplementary Fig. 14.**
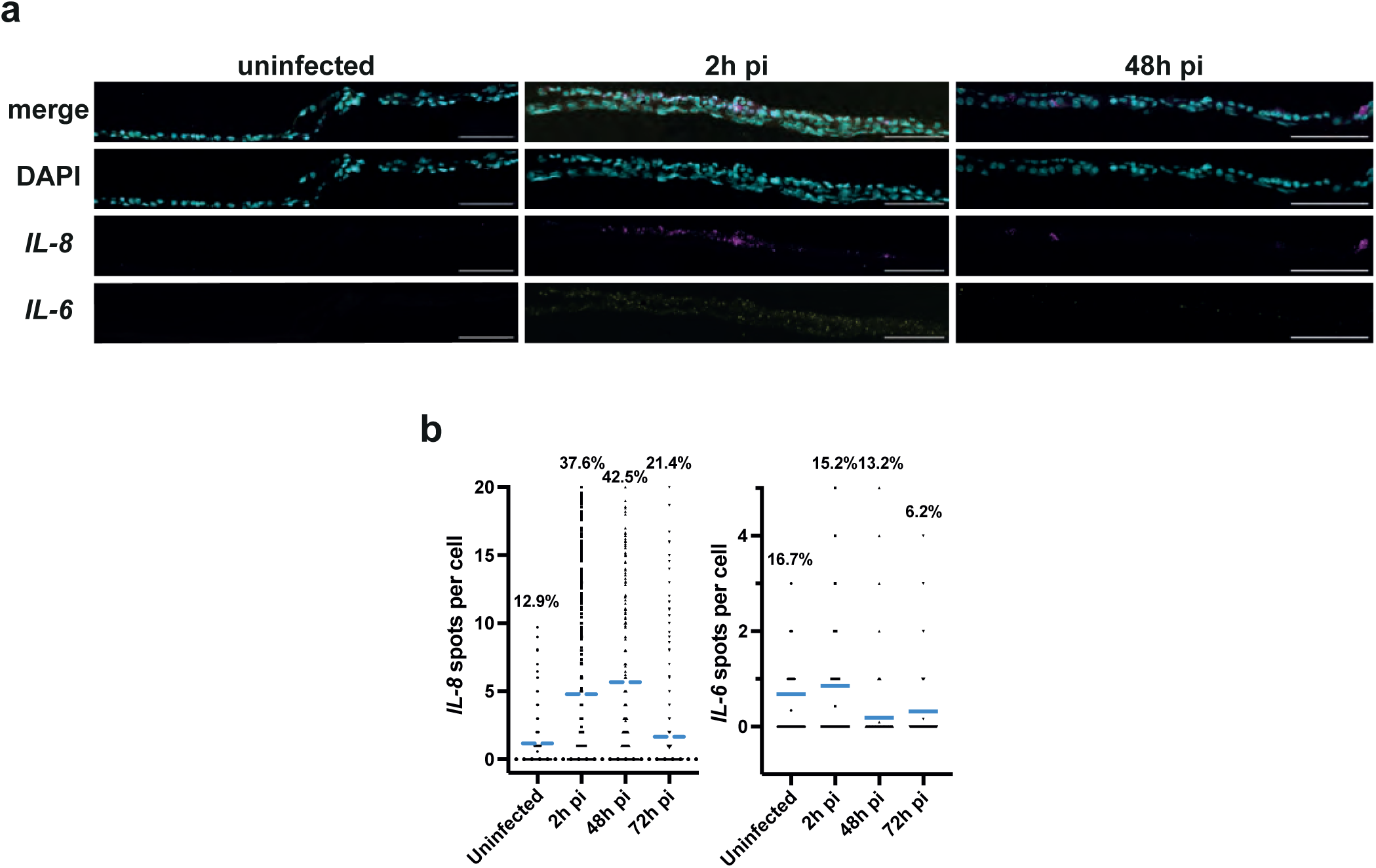
Cytokine expression during *M. tuberculosis* infection of WT-iLoC. **a**, Representative RNAscope images of *IL-6* and *IL-8* expression in iLoC, images showing nuclei (cyan), *IL-6* (yellow) and *IL-8* (magenta). Scalebar, 100 µm. **b**, Quantification of *IL-6* and *IL-8* signals per cell of data displayed in a, percentages represent the number of cells with at least one spot of corresponding transcript; n = >850 quantified cells from n = 1 independent experiment.

**Supplementary Fig. 15.**
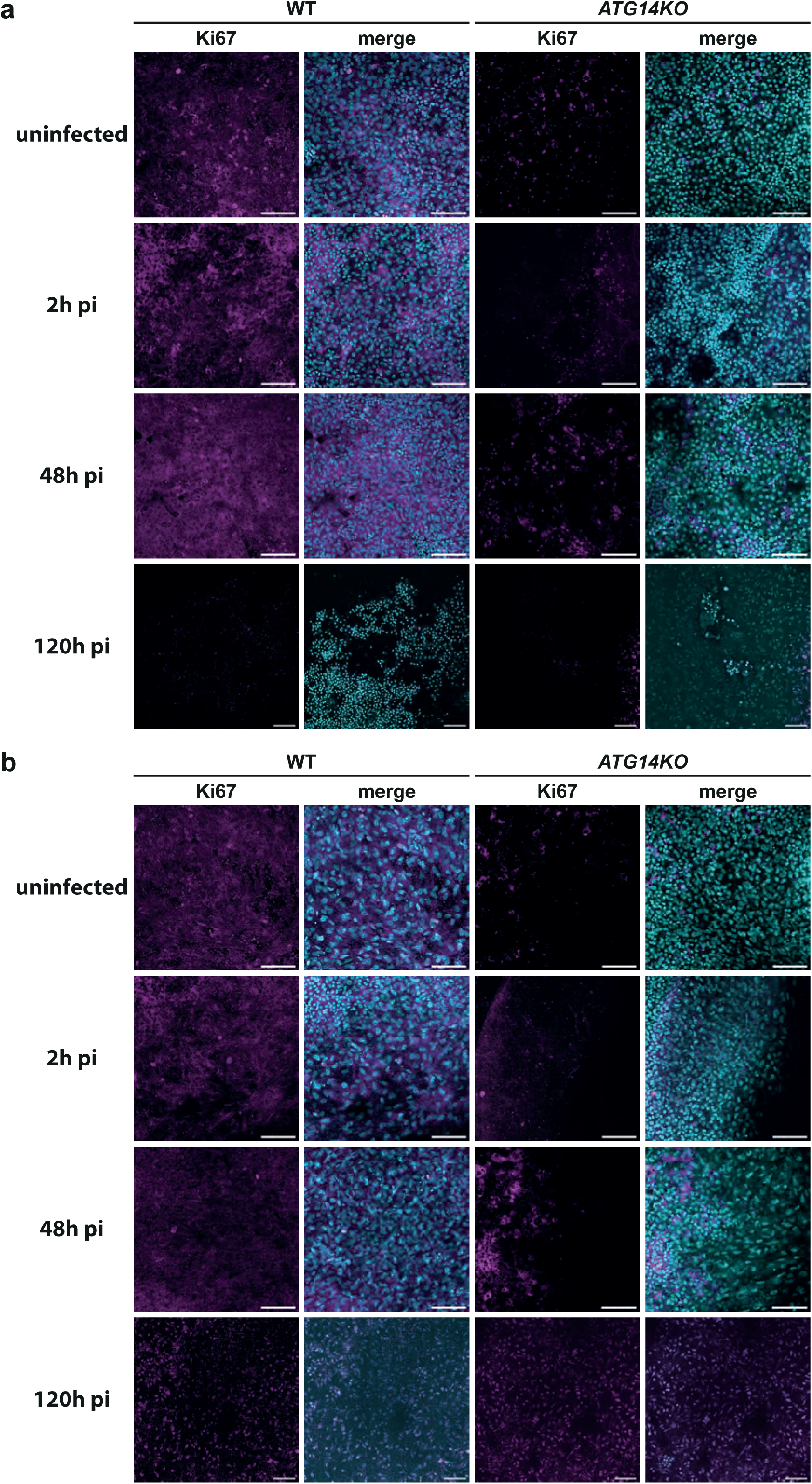
Cell proliferation in the WT-iLoC and *ATG14KO* GE-iLoC after infection with *M. tuberculosis*. **a**, Representative confocal images of cell proliferation in the epithelial side of iLoC or GE-iLoC indicated by Ki67 at different infection timepoints, images showing nuclei (cyan) and Ki67 (magenta). Scalebar, 100 µm. **b**, Representative images of cell proliferation in the endothelial side of iLoC or GE-iLoC indicated by Ki67 at different infection timepoints, images showing nuclei (cyan) and Ki67 (magenta). Scalebar, 100 µm.

**Table S1**. Results from differential expression analysis in each pairwise comparison.

## References

1 Moldoveanu, B. et al. Inflammatory mechanisms in the lung. J Inflamm Res 2, 1–11 (2009).

2 Knudsen, L. & Ochs, M. The micromechanics of lung alveoli: structure and function of surfactant and tissue components. Histochem Cell Biol 150, 661–676 (2018). 10.1007/s00418-018-1747-9

3 Basil, M. C. & Morrisey, E. E. Lung regeneration: a tale of mice and men. Semin Cell Dev Biol 100, 88–100 (2020). 10.1016/j.semcdb.2019.11.006

4 Han, J. J. FDA Modernization Act 2.0 allows for alternatives to animal testing. Artif Organs 47, 449–450 (2023). 10.1111/aor.14503

5 Ingber, D. E. Human organs-on-chips for disease modelling, drug development and personalized medicine. Nat Rev Genet 23, 467–491 (2022). 10.1038/s41576-022-00466-9

6 Si, L. et al. A human-airway-on-a-chip for the rapid identification of candidate antiviral therapeutics and prophylactics. Nat Biomed Eng 5, 815–829 (2021). 10.1038/s41551-021-00718-9

7 Plebani, R. et al. Modeling pulmonary cystic fibrosis in a human lung airway-on-a-chip: Cystic fibrosis airway chip. J Cyst Fibros (2021). 10.1016/j.jcf.2021.10.004

8 Stucki, J. D. et al. Medium throughput breathing human primary cell alveolus-on-chip model. Sci Rep 8, 14359 (2018). 10.1038/s41598-018-32523-x

9 Bai, H. et al. Mechanical control of innate immune responses against viral infection revealed in a human lung alveolus chip. Nat Commun 13, 1928 (2022). 10.1038/s41467-022-29562-4

10 van Riet, S. et al. Organoid-based expansion of patient-derived primary alveolar type 2 cells for establishment of alveolus epithelial Lung-Chip cultures. Am J Physiol Lung Cell Mol Physiol 322, L526–L538 (2022). 10.1152/ajplung.00153.2021

11 Baptista, D. et al. 3D Lung-on-Chip Model Based on Biomimetically Microcurved Culture Membranes. ACS Biomater Sci Eng 8, 2684–2699 (2022). 10.1021/acsbiomaterials.1c01463

12 Huang, D. et al. Reversed-engineered human alveolar lung-on-a-chip model. Proc Natl Acad Sci U S A 118 (2021). 10.1073/pnas.2016146118

13 Huh, D. et al. Reconstituting organ-level lung functions on a chip. Science 328, 1662–1668 (2010). 10.1126/science.1188302

14 Sengupta, A. et al. A New Immortalized Human Alveolar Epithelial Cell Model to Study Lung Injury and Toxicity on a Breathing Lung-On-Chip System. Front Toxicol 4, 840606 (2022). 10.3389/ftox.2022.840606

15 Mishra, R. et al. Mechanopathology of biofilm-like Mycobacterium tuberculosis cords. Cell (2023). 10.1016/j.cell.2023.09.016

16 Thacker, V. V. et al. A lung-on-chip model of early Mycobacterium tuberculosis infection reveals an essential role for alveolar epithelial cells in controlling bacterial growth. Elife 9 (2020). 10.7554/eLife.59961

17 Thacker, V. V. et al. Rapid endotheliitis and vascular damage characterize SARS-CoV-2 infection in a human lung-on-chip model. EMBO Rep 22, e52744 (2021). 10.15252/embr.202152744

18 Burgess, C. L. et al. Generation of human alveolar epithelial type I cells from pluripotent stem cells. Cell Stem Cell (2024). 10.1016/j.stem.2024.03.017

19 Huang, J. et al. SARS-CoV-2 Infection of Pluripotent Stem Cell-Derived Human Lung Alveolar Type 2 Cells Elicits a Rapid Epithelial-Intrinsic Inflammatory Response. Cell Stem Cell 27, 962–973 e967 (2020). 10.1016/j.stem.2020.09.013

20 Jacob, A. et al. Derivation of self-renewing lung alveolar epithelial type II cells from human pluripotent stem cells. Nat Protoc 14, 3303–3332 (2019). 10.1038/s41596-019-0220-0

21 Gotoh, S. et al. Generation of alveolar epithelial spheroids via isolated progenitor cells from human pluripotent stem cells. Stem Cell Reports 3, 394–403 (2014). 10.1016/j.stemcr.2014.07.005

22 Yamamoto, Y. et al. Long-term expansion of alveolar stem cells derived from human iPS cells in organoids. Nat Methods 14, 1097–1106 (2017). 10.1038/nmeth.4448

23 Heo, H. R. et al. Human pluripotent stem cell-derived alveolar epithelial cells are alternatives for in vitro pulmotoxicity assessment. Sci Rep 9, 505 (2019). 10.1038/s41598-018-37193-3

24 Bluhmki, T. et al. Functional human iPSC-derived alveolar-like cells cultured in a miniaturized 96-Transwell air-liquid interface model. Sci Rep 11, 17028 (2021). 10.1038/s41598-021-96565-4

25 Sun, Y. L. et al. Heterogeneity in Human Induced Pluripotent Stem Cell-derived Alveolar Epithelial Type II Cells Revealed with ABCA3/SFTPC Reporters. Am J Respir Cell Mol Biol 65, 442–460 (2021). 10.1165/rcmb.2020-0259OC

26 Ohnishi, Y. et al. Screening of factors inducing alveolar type 1 epithelial cells using human pluripotent stem cells. Stem Cell Reports 19, 529–544 (2024). 10.1016/j.stemcr.2024.02.009

27 Schneider, J. P., Wrede, C. & Muhlfeld, C. The Three-Dimensional Ultrastructure of the Human Alveolar Epithelium Revealed by Focused Ion Beam Electron Microscopy. Int J Mol Sci 21 (2020). 10.3390/ijms21031089

28 Stahlman, M. T., Gray, M. P., Falconieri, M. W., Whitsett, J. A. & Weaver, T. E. Lamellar body formation in normal and surfactant protein B-deficient fetal mice. Lab Invest 80, 395–403 (2000). 10.1038/labinvest.3780044

29 Ochs, M. The closer we look the more we see? Quantitative microscopic analysis of the pulmonary surfactant system. Cell Physiol Biochem 25, 27–40 (2010). 10.1159/000272061

30. Weaver, T. E., Nogee, L. M. & Jobe, A. H. in Fetal and Neonatal Lung Development: Clinical Correlates and Technologies for the Future (eds Alan H. Jobe, Jeffrey A. Whitsett, & Steven H. Abman) 141-163 (Cambridge University Press, 2016).

31 Patsch, C. et al. Generation of vascular endothelial and smooth muscle cells from human pluripotent stem cells. Nat Cell Biol 17, 994–1003 (2015). 10.1038/ncb3205

32 Bernard, E. M. et al. M. tuberculosis infection of human iPSDM reveals complex membrane dynamics during xenophagy evasion. J Cell Sci (2020). 10.1242/jcs.252973

33 Lachmann, N. et al. Large-scale hematopoietic differentiation of human induced pluripotent stem cells provides granulocytes or macrophages for cell replacement therapies. Stem Cell Reports 4, 282–296 (2015). 10.1016/j.stemcr.2015.01.005

34 Litvack, M. L. et al. Alveolar-like Stem Cell-derived Myb(-) Macrophages Promote Recovery and Survival in Airway Disease. Am J Respir Crit Care Med 193, 1219–1229 (2016). 10.1164/rccm.201509-1838OC

35 Mitsi, E. et al. Human alveolar macrophages predominately express combined classical M1 and M2 surface markers in steady state. Respir Res 19, 66 (2018). 10.1186/s12931-018-0777-0

36 Weibel, E. R. On the tricks alveolar epithelial cells play to make a good lung. Am J Respir Crit Care Med 191, 504–513 (2015). 10.1164/rccm.201409-1663OE

37 Eckert, H., Lux, M. & Lachmann, B. The role of alveolar macrophages in surfactant turnover. An experimental study with metabolite VIII of bromhexine (Ambroxol). Lung 161, 213–218 (1983). 10.1007/BF02713866

38 Branchett, W. J. & Lloyd, C. M. Regulatory cytokine function in the respiratory tract. Mucosal Immunol 12, 589–600 (2019). 10.1038/s41385-019-0158-0

39 Bain, C. C. & MacDonald, A. S. The impact of the lung environment on macrophage development, activation and function: diversity in the face of adversity. Mucosal Immunol 15, 223–234 (2022). 10.1038/s41385-021-00480-w

40 Fu, Y. et al. An IL-9-pulmonary macrophage axis defines the allergic lung inflammatory environment. Sci Immunol 7, eabi9768 (2022). 10.1126/sciimmunol.abi9768

41 Rabolli, V. et al. The alarmin IL-1alpha is a master cytokine in acute lung inflammation induced by silica micro- and nanoparticles. Part Fibre Toxicol 11, 69 (2014). 10.1186/s12989-014-0069-x

42 Rafii, S. et al. Platelet-derived SDF-1 primes the pulmonary capillary vascular niche to drive lung alveolar regeneration. Nat Cell Biol 17, 123–136 (2015). 10.1038/ncb3096

43 Sikkema, L. et al. An integrated cell atlas of the lung in health and disease. Nat Med 29, 1563–1577 (2023). 10.1038/s41591-023-02327-2

44 Pahari, S. et al. A new tractable method for generating human alveolar macrophage-like cells in vitro to study lung inflammatory processes and diseases. mBio 14, e0083423 (2023). 10.1128/mbio.00834-23

45 Hernandez-Pando, R. et al. Persistence of DNA from Mycobacterium tuberculosis in superficially normal lung tissue during latent infection. Lancet 356, 2133–2138 (2000). 10.1016/s0140-6736(00)03493-0

46 Aylan, B. et al. ATG7 and ATG14 restrict cytosolic and phagosomal Mycobacterium tuberculosis replication in human macrophages. Nat Microbiol 8, 803–818 (2023). 10.1038/s41564-023-01335-9

47 Nawroth, J. C. et al. Stem cell-based Lung-on-Chips: The best of both worlds? Adv Drug Deliv Rev 140, 12–32 (2019). 10.1016/j.addr.2018.07.005

48 Pai, M. et al. Tuberculosis. Nat Rev Dis Primers **2**, 16076 (2016). 10.1038/nrdp.2016.76

49 Rothchild, A. C. et al. Alveolar macrophages generate a noncanonical NRF2-driven transcriptional response to Mycobacterium tuberculosis in vivo. Sci Immunol 4 (2019). 10.1126/sciimmunol.aaw6693

50 Goodwin, A. M. In vitro assays of angiogenesis for assessment of angiogenic and anti-angiogenic agents. Microvasc Res 74, 172–183 (2007). 10.1016/j.mvr.2007.05.006

51 Stringer, C., Wang, T., Michaelos, M. & Pachitariu, M. Cellpose: a generalist algorithm for cellular segmentation. Nat Methods 18, 100–106 (2021). 10.1038/s41592-020-01018-x

52 Ulicna, K., Vallardi, G., Charras, G. & Lowe, A. R. Automated Deep Lineage Tree Analysis Using a Bayesian Single Cell Tracking Approach. Frontiers in Computer Science 3 (2021). 10.3389/fcomp.2021.734559

53 Sofroniew, N., et al. napari: a multi-dimensional image viewer for Python. Zenodo (2022).

54 Stuart, T. et al. Comprehensive Integration of Single-Cell Data. Cell 177, 1888–1902 e1821 (2019). 10.1016/j.cell.2019.05.031

55 Butler, A., Hoffman, P., Smibert, P., Papalexi, E. & Satija, R. Integrating single-cell transcriptomic data across different conditions, technologies, and species. Nat Biotechnol 36, 411–420 (2018). 10.1038/nbt.4096

56 Fu, R. et al. clustifyr: an R package for automated single-cell RNA sequencing cluster classification. F1000Res **9**, 223 (2020). 10.12688/f1000research.22969.2

57 Subramanian, A. et al. Gene set enrichment analysis: a knowledge-based approach for interpreting genome-wide expression profiles. Proc Natl Acad Sci U S A 102, 15545–15550 (2005). 10.1073/pnas.0506580102

58 Wu, T. et al. clusterProfiler 4.0: A universal enrichment tool for interpreting omics data. Innovation (Camb) 2, 100141 (2021). 10.1016/j.xinn.2021.100141

59 Liberzon, A. et al. Molecular signatures database (MSigDB) 3.0. Bioinformatics 27, 1739–1740 (2011). 10.1093/bioinformatics/btr260

